# RNA polymerase II-TFIIE-TFIIH interface functions in transcription start site selection in *Saccharomyces cerevisiae*

**DOI:** 10.1101/2025.11.05.686797

**Authors:** Pratik Basnet, Yunye Zhu, Irina O. Vvedenskaya, Payal Arora, Staci Hammer, Brittany McVicar, Shawn Alex, Bryce E. Nickels, Craig D. Kaplan

## Abstract

During transcription initiation in *Saccharomyces cerevisiae*, RNA polymerase II (Pol II) and general transcription factors (GTFs) assemble upstream of transcription start sites (TSSs) to form the pre-initiation complex (PIC). In this model organism, yeast, the PIC selects TSSs through a unidirectional scanning mechanism referred to as promoter scanning. Previous studies have shown that the TFIIH subunit Tfb3 connects TFIIH to the rest of the PIC through interactions with Pol II and the GTF TFIIE. Activities within the PIC that influence TSS selection can do so by control of initiation efficiency at individual TSSs or by control of TSS scanning (either rate of scanning or scanning processivity). To understand how this critical interface withing the PIC participates in scanning, we used genetic screens to identify *tfb3* and *tfa1* mutants that alter initiation using initiation-linked phenotypes. We found mutations within the TFIIH-Pol II-TFIIE interface able to alter promoter scanning in either upstream or downstream directions, suggesting that changes to this interface can fine-tune scanning. Subsets of alleles were analyzed using TSS sequencing approaches, showing that tested *tfb3* and *tfa1* alleles shift TSS distributions across most genomic promoters. Genetic interaction and genomic analysis revealed that the Tfb3 interfaces with Rpb7 and Tfa1 separately contribute to promoter scanning, and that *tfb3* alleles exhibit additive effects with scanning processivity mutants in, consistent with Tfb3-PIC interactions modulating scanning processivity. The ability of this interface to easily modulate scanning in both directions is consistent with the types of changes that might incrementally allow promoter scanning to have evolved.

## INTRODUCTION

In eukaryotes, RNA polymerase II (Pol II) transcribes all protein-coding genes and numerous noncoding RNAs in the first step of gene expression. This process proceeds through three consecutive steps: initiation, elongation, and termination (ROEDER 2019). The first step in transcription, initiation, requires promoter opening, insertion of the template DNA strand into the Pol II active site, and transcription start site (TSS) selection (HAHN 2004; CRAMER *et al*. 2008; HABERLE AND STARK 2018). Most promoters have multiple TSSs, and the precise choice of TSS can influence mRNA content and gene expression (CARNINCI *et al*. 2005; ZHANG AND DIETRICH 2005; KAPLAN 2013; NEPAL *et al*. 2013; HABERLE *et al*. 2014). Initiation is a highly regulated process and requires the activities of RNA Pol II along with several general transcription factors (GTFs) (CONAWAY AND CONAWAY 1993; KORNBERG 2007; GRÜNBERG AND HAHN 2013). Pol II and GTFs are recruited and assembled upstream of the TSS on promoter DNA to form a pre-initiation complex (PIC).

Although PIC components and their structures are highly conserved (BURATOWSKI *et al*. 1988; TOMKO *et al*. 2021), TSS selection in most eukaryotes is distinct from that in budding yeast. In higher eukaryotes, transcription initiation for promoters containing a TATA-box core promoter element occurs at ∼30-31 bp downstream of the TATA-box (KADONAGA 2012; DANINO *et al*. 2015; DUDNYK *et al*. 2024). In contrast, initiation in *S. cerevisiae* occurs at multiple TSSs ranging from ∼40 to 120 bp downstream of a TATA-box when present (STRUHL 1989; ZHANG AND DIETRICH 2005). The mechanism of TSS selection in yeast involves a unidirectional scanning mechanism wherein the PIC surveys downstream positions for suitable TSSs to initiate transcription. This mechanism of initiation in yeast is known as “promoter scanning” (GIARDINA AND LIS 1993; HAMPSEY 2006; KUEHNER AND BROW 2006; FAZAL *et al*. 2015; MURAKAMI *et al*. 2015). Genetic and genomic studies have established promoter scanning as a general feature of yeast transcription initiation(QIU *et al*. 2020; LU AND LIN 2021; YANG *et al*. 2022; ZHU *et al*. 2024).

The promoter scanning model posits that individual TSSs possess intrinsic differences in their likelihood of being used, and genetic and molecular data indicate that changes in the activity of Pol II and GTFs can shift TSS usage upstream or downstream, as reflected by changes to the overall distribution of TSS usage at promoters (PINTO *et al*. 1992; SUN 1996; BANGUR *et al*. 1997; WU *et al*. 1999; FAITAR *et al*. 2001; GHAZY *et al*. 2004; FREIRE-PICOS 2005; KAPLAN *et al*. 2012; BRABERG *et al*. 2013; JIN AND KAPLAN 2015; ZHAO *et al*. 2021). Promoter scanning has been proposed to be regulated through two distinct networks, the first relating to the innate initiation efficiency of individual TSSs and Pol II catalytic activity, and the second relating to TFIIH DNA translocase activity as the proposed engine for promoter scanning (ZHAO *et al*. 2021; ZHU *et al*. 2024; ARORA *et al*. 2025). While TSS selection is primarily dictated by the activities of TFIIH and Pol II, there are multiple interactions among PIC components during initiation. How each PIC component and their interactions influence the TSS selection mechanism, either through communication to different enzymatic activities in the PIC or through structural means, is not clear at the molecular level.

TFIIH, a multiprotein complex of 11 subunits in yeast, functions to drive promoter opening for Pol II promoters in eukaryotes (SVEJSTRUP *et al*. 1994; RANISH *et al*. 2004; MURAKAMI *et al*. 2012; GRÜNBERG AND HAHN 2013). To do this, TFIIH translocates downstream DNA towards the PIC using the ATPase activity of its Ssl2 subunit (the yeast homolog of human XPB) (FAZAL *et al*. 2015; FISHBURN *et al*. 2015; FISHBURN *et al*. 2016; TOMKO *et al*. 2021). During promoter scanning initiation, TFIIH has been proposed to continue translocating on downstream DNA subsequent to promoter opening, allowing downstream TSSs to be accessed by the PIC (FAZAL *et al*. 2015; TOMKO *et al*. 2021; ZHAO *et al*. 2021; BASSETT *et al*. 2022). Structural studies have shown that the TFIIH component Tfb3, the yeast homolog of human MAT1, links TFIIH to the rest of the PIC via interactions with the Pol II subunit Rpb7 and TFIIE, through Tfa1 (CHEN *et al*. 2021; SCHILBACH *et al*. 2021). Intriguingly, yeast PICs lacking the TFIIK module (Tfb3, Kin28, Ccl1) or just the Tfb3 RING domain initiate far upstream in vitro near the position observed for TSS selection in other eukaryotes (MURAKAMI *et al*. 2015; YANG *et al*. 2022). This work highlighted the potential importance of these interactions in the evolution of promoter scanning.

Genetic and molecular data suggest that Tfb3-Rpb7 interactions are functionally important for promoter scanning in vivo, as mutations on either side of this interface cause upstream shifts to TSS usage (BRABERG *et al*. 2013; YANG *et al*. 2022; YANG *et al*. 2024). Given the strong effects of Tfb3 on initiation in vitro and in vivo, the interface between Pol II and TFIIH has been suggested as one facet for how promoter scanning has evolved in yeast, with residues differing between yeast and human located within Tfb3 shown to affect TSS selection in vivo (YANG *et al*. 2024). One residue in particular, Tfb3 P51 appears to work by supporting adjacent Tfb3 residues in electrostatic interaction with Rpb7 (YANG *et al*. 2024). Interestingly, the MAT1 (human Tfb3)-Rpb7 interface may also undergo remodeling during human promoter opening (AIBARA *et al*. 2021). In reconstituted closed complex PIC structures, MAT1-Rpb7 interactions are intact, including through conserved residues that can support salt bridges in both human and yeast PICs. Putatively, upon XPB translocation this interface breaks and subsequently density for the MAT1 RING domain is not observed, suggesting these interactions may control coupling of Ssl2/XPB to the rest of the PIC during promoter opening/promoter scanning (AIBARA *et al*. 2021).

Here, we have undertaken genetic and genomic analysis of the Pol II-TFIIH-TFIIE interface functions in TSS selection. We have identified novel *tfb3* and *tfa1* alleles by genetic screens for transcription-related phenotypes. For both Tfb3 and Tfa1, we identified two classes of alleles that produce distinct polarity of shifts in TSS usage in vivo. In addition, genetic interactions of *tfb3* alleles with either *ssl2* alleles or *sub1*Δ were primarily additive, suggesting that Tfb3 defects can alter TFIIH processivity. We have probed the contributions of electrostatic interactions between Rpb7 and Tfb3 in promoter scanning, providing evidence for the contribution of two conserved salt bridges at this interface for TSS selection. Intramolecular and intermolecular genetic interactions support that contacts between both Tfb3-Rpb7 and Tfb3-Tfa1 additively contribute to promoter scanning. Importantly, intramolecular suppression of putative interface disrupting mutations that shift TSSs upstream by opposite polarity, downstream-shifting mutations are consistent with the latter class enhancing interactions at the interface. In parallel, we mapped TSSs genome-wide for representative *tfb3* and *tfa1* alleles, which demonstrated corresponding global upstream or downstream shifts in TSS usage. Collectively, our findings indicate that the Pol II-TFIIH-TFIIE interface, mediated through Tfb3-Rpb7 and Tfb3-Tfa1 interactions, plays a critical role in the promoter scanning mechanism and can be tuned by mutation.

## RESULTS

### Genetic screens for *tfb3* and *tfa1* alleles show transcription-related phenotypes

To dissect how promoter scanning governs TSSs selection during transcription initiation in yeast and to probe the role of an RNA Pol II-GTF interface, we performed genetic screens for *tfb3* and *tfa1* alleles with transcription-initiation-related phenotypes (see Materials and methods). We applied a battery of phenotypes during our genetic screening. Two of these phenotypes are related to transcription initiation at the *IMD2* promoter (**Figure 1a**) (KAPLAN *et al*. 2012; BRABERG *et al*. 2013). One initiation phenotype reflects the sensitivity of yeast cells to Inosine-5′-monophosphate dehydrogenase (IMPDH) inhibitors such as the drug mycophenolic acid (MPA) (SHAW AND REINES 2000; HYLE *et al*. 2003). IMDPH is the rate limiting step in GTP synthesis, and *IMD2* encodes an MPA-resistant form of IMDPH and is necessary for cell viability in the presence of MPA. When MPA is present, the reduction in GTP levels leads to usage of a downstream TSS that allows for production of *IMD2* mRNA (SHAW AND REINES 2000; HYLE *et al*. 2003; JENKS *et al*. 2008; KUEHNER AND BROW 2008; KWAPISZ *et al*. 2008). Mutants that shift TSSs upstream reduce the usage of the downstream TSS required for MPA resistance and are therefore sensitive to MPA (**Figure 1a**, **top**) (KAPLAN *et al*. 2012; JIN AND KAPLAN 2015; ZHAO *et al*. 2021). We screened for downstream-shifting mutants using the converse *IMD2* phenotype: the constitutive usage of the downstream *IMD2* TSS seen for a wide range of initiation mutants. We detect constitutive *IMD2* expression using the reporter allele *imd2*Δ*::HIS3*, where *HIS3* replaces the *IMD2* ORF as it can easily be observed as a His^+^ phenotype (**Figure 1a**, **bottom**) (MALIK *et al*. 2017; ZHAO *et al*. 2021). These phenotypes have been used as genetic tools to probe initiation defects in previous studies relating to Pol II and other GTF mutants.

**Figure 1:**
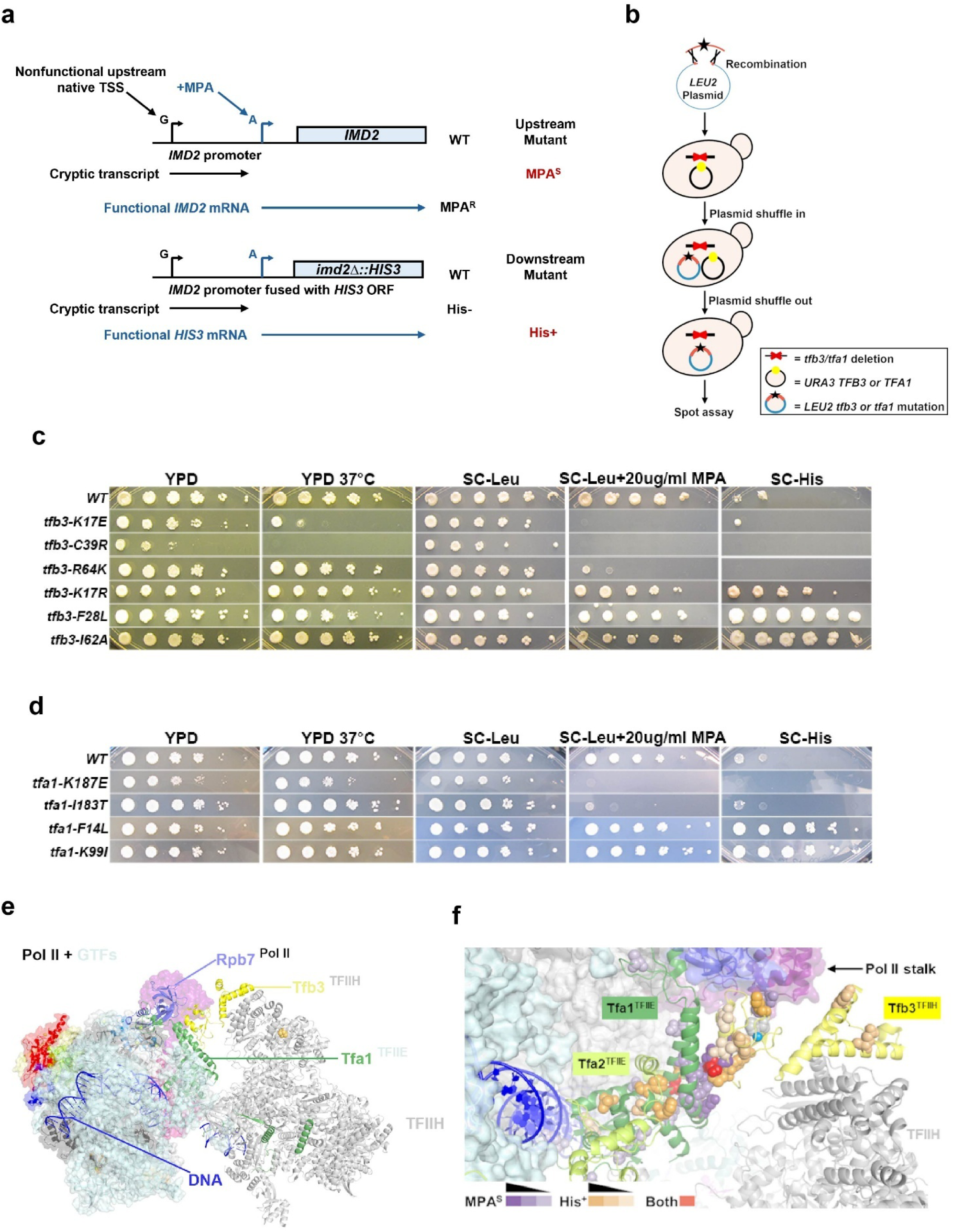
Genetic screens identify novel mutants with initiation phenotypes at the interface between Pol II/TFIIE and TFIIH. **a.** Schematic representation of two transcriptional phenotypes used in this study. Mycophenolic acid (MPA) sensitivity: MPA inhibits the yeast inosine monophosphate dehydrogenase (IMPDH) enzymes encoded by *IMD3* and *IMD4*, leading to GTP depletion. In response, wild-type cells induce *IMD2* expression by a TSS shift from an upstream “G” TSSs, which produces nonfunctional cryptic transcripts, to a downstream “A” TSS that yields a functional *IMD2* mRNA. Mutants that are unable to shift the usage to the downstream TSS show MPA^S^ (top). Suppression of *imd2*Δ*::HIS3*: The *IMD2* ORF was replaced with the *HIS3* to generate the reporter construct. In the absence of MPA, wild-type cells initiate transcription from upstream TSSs of the *IMD2* promoter, producing nonfunctional transcripts and failing to grow on medium lacking histidine (SC-His). Mutants that shift transcription constitutively to the downstream TSS express *HIS3* and show a His⁺ phenotype (bottom). **b.** Schematic of genetic screening by PCR-based random mutagenesis coupled with gap repair/plasmid shuffle to examine *tfb3* or *tfa1* alleles. **c.** Initiation phenotypes of select representative novel *tfb3* mutants obtained from genetic screening. **d.** Initiation phenotypes of select representative novel *tfa1* mutants obtained from genetic screening. 10-fold serial dilutions of saturated cultures of WT and mutant strains were plated on different phenotyping media. **e.** Positions of Tfb3 and Tfa1 relative to TFIIH and Pol II in the yeast PIC (PDB 7O4J) (SCHILBACH *et al*. 2021). **f.** Mapping novel *tfb3 and tfa1* mutant residues shown as spheres on the cartoon representation of Tfb3 and Tfa1 protein structures from (**e**). Residues are colored according to phenotype: shades of purple indicate increasing MPA^S^ mutant strength; shades of orange indicate increasing His⁺ mutant strength; exhibiting with distinct substitutions conferring MPA^S^ or His⁺ phenotypes are shown in red.

We focused on *TFA1*, encoding a component of TFIIE, and *TFB3*, encoding a component of TFIIH because of the structurally described Rpb7-Tfb3-Tfa1 interface in the yeast PIC (SCHILBACH *et al*. 2017; SCHILBACH *et al*. 2021). Prior functional work implicated the RING finger domain of Tfb3 as critical for proper TSS selection during scanning. Tfb3 binds in a pocket formed by the Tfa1 E-linker helices and the OB domain of Rpb7 (SCHILBACH *et al*. 2017; YANG *et al*. 2022). We performed *TFB3* and *TFA1* genetic screens by a gap-repair strategy using random PCR mutagenesis followed by plasmid shuffling (**Figure 1b**) (**Materials and methods**). We identified 31 *tfb3* and 26 *tfa1* alleles each containing a unique single amino acid substitution (**Table 1 and Table 2**). Mutant growth phenotypes are shown in spot assays. We identified alleles for both *TFA1* and *TFB3* with presumptive TSS-shifting phenotypes of each polarity, MPA^s^ presumptive upstream shifting and His^+^ presumptive downstream shifting alleles (**Figures 1c**, **1d, S1a, S1b**). We failed to identify Spt^-^ mutants for either *TFB3* or *TFA1* screen, which is distinct from a previous genetic screen where these were identified as a subset of *ssl2* His^+^ alleles in Zhao et al (ZHAO *et al*. 2021). Additionally, a subset of these novel *tfb3 and tfa1* mutants showed Gal^R^ phenotypes from the suppression of *gal10*Δ*56*, a *GAL10* allele that is sensitive to transcription defects (GREGER 1998; GREGER *et al*. 2000; KAPLAN *et al*. 2005). These Gal^R^ phenotypes were only exhibited by a fraction of MPA^s^ alleles, for both *tfb3* (*C13S*, *C16R*, *S23P* and *I183F*) and *tfa1* (*K187E, K187N* and *I183F*) (**Figures S1a, S1b**).

**Table 1:** *tfb3* alleles with single amino acid substitutions (Ts = temperature sensitive and, Cs = cold sensitive phenotypes).

| # | Substitution position | Amino acid change | Phenotypes |
| --- | --- | --- | --- |
| 1 | 7 | E7K | His<sup>+</sup> |
| 2 | 12 | M12K | His<sup>+</sup> |
| 3 | 13 | C13S | MPA<sup>S</sup>/Gal<sup>R</sup>/Ts |
| 4 | 15 | I15N | His<sup>+</sup> |
| 5 | 16 | C16R | MPA<sup>S</sup>/Gal<sup>R</sup>/Ts/Cs |
| 6 | 17 | K17E | MPA<sup>S</sup>/Ts |
| 7 | 17 | K17R | His<sup>+</sup> |
| 8 | 21 | Y21H | MPA<sup>S</sup> |
| 9 | 22 | L22P | MPA<sup>S</sup>/Gal<sup>R</sup>/Ts/Cs |
| 10 | 23 | S23P | MPA<sup>S</sup>/Gal<sup>R</sup>/Ts/Cs |
| 11 | 28 | F28L | His<sup>+</sup> |
| 12 | 38 | I38F | His<sup>+</sup> |
| 13 | 39 | C39R | MPA<sup>S</sup>/Ts/Cs |
| 14 | 44 | D44G | His<sup>+</sup> |
| 15 | 46 | I46T | His<sup>+</sup> |
| 16 | 56 | Y56C | His<sup>+</sup> |
| 17 | 56 | Y56F | His<sup>+</sup> |
| 18 | 62 | I62A | His<sup>+</sup> |
| 19 | 62 | I62T | His<sup>+</sup> |
| 20 | 64 | R64K | MPA<sup>S</sup> |
| 21 | 66 | N66D | MPA<sup>S</sup> |
| 22 | 68 | F68L | MPA<sup>S</sup> |
| 23 | 68 | F68S | Gal<sup>R</sup> |
| 24 | 68 | F68V | MPA<sup>S</sup> |
| 25 | 73 | F73S | MPA<sup>S</sup>/Gal<sup>R</sup> |
| 26 | 88 | V88E | Gal<sup>R</sup> |
| 27 | 112 | E112K | His<sup>+</sup> |
| 28 | 112 | E112V | His<sup>+</sup> |
| 29 | 131 | E131V | His<sup>+</sup> |
| 30 | 204 | D204N | His<sup>+</sup> |
| 31 | 218 | L218S | MPA<sup>S</sup> |

**Table 2:** *tfa1* alleles with single amino acid substitutions (Ts = temperature sensitive and, Cs = cold sensitive phenotypes).

| # | Substitution position | Amino acid change | Phenotypes |
| --- | --- | --- | --- |
| 1 | 12 | L12H | MPA<sup>S</sup> |
| 2 | 14 | F14L | His<sup>+</sup> |
| 3 | 17 | V17D | His<sup>+</sup>/Ts |
| 4 | 20 | F20S | His<sup>+</sup> |
| 5 | 29 | L29P | MPA<sup>S</sup> |
| 6 | 33 | L33P | MPA<sup>S</sup> |
| 7 | 35 | H35R | MPA<sup>S</sup> |
| 8 | 57 | L57P | His<sup>+</sup> |
| 9 | 64 | D64G | MPA<sup>S</sup> |
| 10 | 80 | K80E | MPA<sup>S</sup> |
| 11 | 99 | K99I | His<sup>+</sup> |
| 12 | 99 | K99R | MPA<sup>S</sup> |
| 13 | 103 | H103R | His<sup>+</sup> |
| 14 | 106 | V106L | MPA<sup>S</sup>/Ts |
| 15 | 124 | C124S | MPA<sup>S</sup>/Ts |
| 16 | 176 | M176T | MPA<sup>S</sup> |
| 17 | 183 | I183T | MPA<sup>S</sup> |
| 18 | 183 | I183F | MPA<sup>S</sup> |
| 19 | 183 | I183V | MPA<sup>S</sup> |
| 20 | 186 | L186S | MPA<sup>S</sup> |
| 21 | 187 | K187E | MPA<sup>S</sup>/Gal<sup>R</sup> |
| 22 | 187 | K187N | MPA<sup>S</sup>/Gal<sup>R</sup> |
| 23 | 187 | K187Q | MPA<sup>S</sup>/Gal<sup>R</sup> |
| 24 | 190 | D190G | His<sup>+</sup> |
| 25 | 190 | D190N | His<sup>+</sup> |
| 26 | 250 | R250G | His<sup>+</sup> |

### *tfb3* and *tfa1* MPA^S^ alleles map to the TFIIE-TFIIH interface

Mapping substituted residues of *tfb3* and *tfa1* alleles onto the structure of the yeast Pol II PIC revealed that MPA^s^ mutants were clustered phenotype-selectively at the Tfb3-Tfa1 interface, the Tfb3-Rpb7 interface, and in residues important for structural integrity (Zn binding) in the Tfb3 RING domain (SCHILBACH *et al*. 2017; SCHILBACH *et al*. 2021) (**Figures 1e**, **1f**). These results are in line with collaborative studies from our lab and the Murakami lab showing that Tfb3 interactions with Rpb7 are critical for promoter scanning and usage of downstream TSSs (YANG *et al*. 2022). Our screen extends these results and is consistent with disruptions of structural integrity across Tfb3 PIC-interfaces causing MPA sensitivity. *tfb3* alleles at either the Tfb3-Rpb7 interface (*e.g. tfb3-R64K*, *tfb3-N66D*) or Tfb3-Tfa1 interface (e.g. *tfb3-K17E*, *tfb3-Y21H*, *tfb3-L22P*, and *tfb3-S23P*) were the strongest MPA^S^ mutants with a mix of strong and medium MPAs alleles in Tfa1 on the Tfa1 side of the Tfa1-Tfb3 interface (**Figures 1e**, **1f**). Consistent with this interface being important, we identified alleles on both sides of specific interaction between Tfa1 K99 and Tfb3 Y21 (*tfa1-K99I* and *tfb3-Y21H*) and Tfa1 M176 and Tfb3 K17 (*tfa1-M176T* and *tfb3-K17E*).

### His^+^ alleles are represented by more subtle changes and may reflect altered stability or protein dynamics

In contrast to MPA^S^ alleles clearly disrupting protein interactions or key structural components, His^+^ alleles were more distributed with most located at intramolecular or core interactions, though a few alleles mapped to protein interfaces. A general trend for these His⁺ mutants in both Tfa1 and Tfb3 is that substitutions within the protein core typically involved hydrophobic residues replaced by amino acids of similar nonpolar character, preserving overall hydrophobicity. In contrast, His^+^ surface substitutions tended to occur at polar residues, where changes in side-chain length or charge may help maintain or remodel intermolecular interactions.

For Tfb3, most His⁺ alleles were clustered within the RING domain. A recent structural model of the human PIC proposed that early in initiation, remodeling of MAT1/Tfb3 disrupts Pol II-TFIIH coupling by displacing the RING domain, thereby limiting further promoter opening in that system as opposed to the scanning that can occur in yeast (AIBARA *et al*. 2021). The 16 substitutions at 13 residues conferring the His⁺ phenotype in Tfb3 were mostly located at internal positions within the RING domain rather than at intermolecular interfaces or Zn^2+^-binding residues, suggesting that these changes may subtly stabilize or alter the dynamics of the RING fold. For example, residues I38 (I38F) and F28 (F28L) reside in the β1 and β2 sheets that contribute to the characteristic αββαβ architecture of the RING finger domain (Gervais et al., 2001). Likewise, I15 (I15N) and M12 (M12K) lie adjacent to the zinc-binding residues C16 and C13 respectively and confer the opposite phenotype to mutations in zinc-binding cysteines. For Tfa1, nine substitutions across eight residues confer the His⁺ phenotype. Only K99I and H103R are near the Tfa1-Tfb3 interface, while the remaining substitutions are concentrated internally (**Figure 1f**). This distribution suggests that most of these His⁺ alleles primarily affect intramolecular stability rather than disrupting intermolecular contacts.

For both Tfa1 and Tfb3, we identified residues where distinct substitutions at the same position confer either the His⁺ or MPA^S^ phenotypes, arguing for their distinct characters. In Tfb3, the K17 and S23 residues illustrate this dual behavior. The K17E substitution produces a severe MPA^S^ phenotype, likely by disrupting (i) intramolecular interactions involving nearby polar residues within the α-helix or (ii) intermolecular contacts with the Tfa1 interface. Consistent with this, K17 in Tfb3 is positioned to interact with M176 of Tfa1, and the M176T substitution in Tfa1 also yields an MPA^S^ phenotype. In contrast, the K17R substitution, which retains the positive charge but alters side chain character, results in a moderate His⁺ phenotype, suggesting that preservation of electrostatic interactions at this site contribute to the altered phenotype. Similarly, for Tfb3 S23, which is near the Tfa1K99 and Tfb3 Y21 interaction (described below), we identified *tfb3* S23P which shows strong MPA^S^ along with Gal^R^ phenotype, likely by disturbing intermolecular contacts between Tfa1-Tfb3 interface. In contrast, Arora et al. 2025 isolated *tfb3-*S23F alleles with the His^+^ phenotype, supporting idea that this residue is also placed uniquely at Tfb3-Tfa1 interface where depending upon the substation it can either reinforce or disturb the Tfa1-Tfb3 interaction, which is crucial for promoter scanning (ARORA *et al*. 2025).

A parallel pattern was observed for Tfa1 at K99, which forms a hydrogen bond with Y21 of Tfb3 (SCHILBACH *et al*. 2021). The K99R substitution leads to an MPA^S^ phenotype (**Figure S1b**), likely for reasons similar to those observed for Tfb3 K17E, where perturbation of interfacial interactions compromises functional coupling (discussed below). Consistently, the Y21H substitution in Tfb3 also causes an MPA^S^ phenotype (**Figure S1a**). Whereas the K99I substitution in Tfa1 produces a strong His⁺ phenotype (**Figures 1d, S1b**). Given that the K99 residue occupies a unique position at the boundary of alleles associated with MPA^S^ and His⁺ phenotypes, implicating K99 in Tfb3-Tfa1 interface interactions and potentially intramolecular stabilization within Tfa1.

### Novel *tfb3* and *tfa1* alleles display unique patterns of TSS usage, differing from the previously defined ‘efficiency’ and ‘processivity’ mutants

The core promoter of *ADH1* has been used as a model to quantitatively determine TSS usage defects. Previously, we found a perfect concordance between PIC mutants conferring initiation phenotypes at *IMD2* with those that also show quantitative defects at *ADH1* (KAPLAN *et al*. 2012; JIN AND KAPLAN 2015; MALIK *et al*. 2017; ZHAO *et al*. 2021). Furthermore, mutations showing growth phenotypes at *IMD2* and TSS phenotypes at *ADH1* also showed global TSS selection effects across yeast promoters for the subset that have been evaluated (QIU *et al*. 2020; ZHAO *et al*. 2021). We employ primer extension (PE) to measure the TSS usage at *ADH1*, which contains two major TSSs at - 37 and -27 positions relative to the first base of the translation start codon (+1) along with multiple minor TSSs (**Figure 2a**) (HULL *et al*. 1995; SUN 1996). The quantification scheme for observed *ADH1* TSS usage is described in (JIN AND KAPLAN 2015) (**Figure 2b**). Quantifications of PE for *tfb3-L218S* (a putative upstream shifting mutant) and *tfb3-F28L* (a putative downstream shifting mutant) are shown as examples. Consistent with previously studied Pol II and other GTF mutants’ examined by primer extension, all tested *tfb3* and *tfa1* MPA^S^ alleles shifted TSS usage upstream at *ADH1*, whereas tested His^+^ alleles shifted TSS usage downstream (**Figures 2c**, **2d**) (KAPLAN *et al*. 2012; JIN AND KAPLAN 2015; MALIK *et al*. 2017; QIU *et al*. 2020; ZHAO *et al*. 2021). The effects of strong MPA^S^ *tfa1* and *tfb3* alleles appeared intermediate between those previously observed for tested Pol II MPA^S^ alleles and tested *ssl2* MPA^S^ alleles. Pol II alleles have been interpreted as solely acting through increased catalysis, which allows increased efficiency and therefore increases in absolute as well as relative usage of upstream TSSs, whereas *ssl2* alleles were interpreted as being defective in scanning processivity, meaning that primarily downstream TSS usage was truncated, with generally only a relative increase in usage of upstream TSSs (KAPLAN *et al*. 2012; MALIK *et al*. 2017; QIU *et al*. 2020; ZHAO *et al*. 2021). Interestingly, both *tfb3* and *tfa1* MPA^S^ alleles, consistent with their relative phenotypic strengths, simultaneously increase the usage of upstream TSSs while reducing usage of downstream TSSs. As expected, and consistent with their phenotypic strengths, both *tfb3* and *tfa1* His⁺ alleles shifted the TSS distribution downstream by increasing usage of downstream TSSs (see increase in quantification bins 5 and 6) (**Figures 2c**, **2d**).

**Figure 2:**
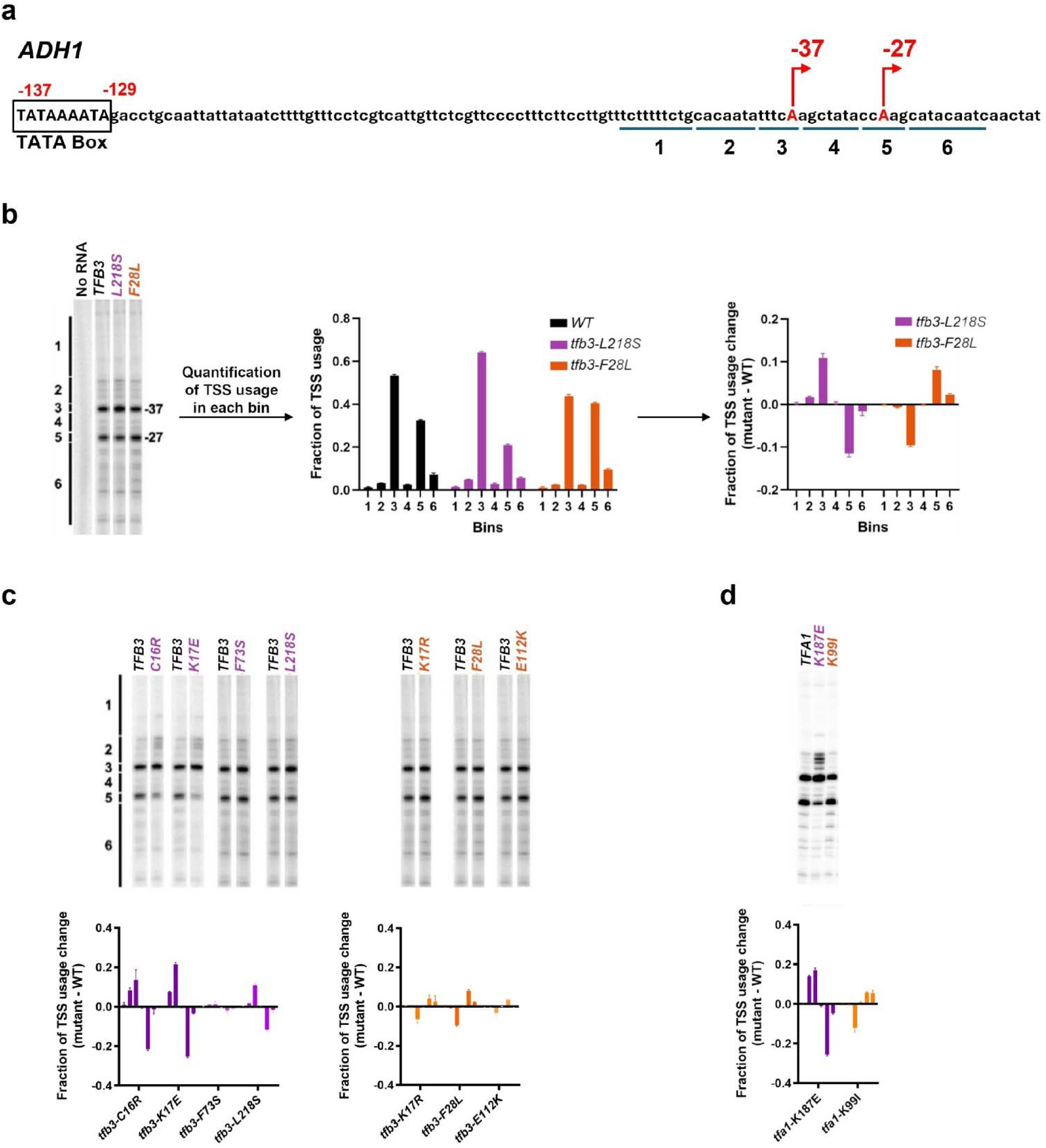
*tfb3* and *tfa1* alleles show TSS distribution alterations at *ADH1*. **a.** Schematic of the TSS region at *ADH1*. The *ADH1* promoter contains two major TSSs at −37 and −27 nucleotides upstream of the *ADH1* start codon (designated as −37 and −27). **b.** Quantification scheme for *ADH1* TSS usage for *tfb3* MPA^S^ and His^+^ mutants. *ADH1* TSS signals were divided into six bins, quantified, and normalized to the total signal from all bins, with the background removed based on “no RNA” reaction signals. The change in the TSS usage fraction for mutants relative to WT was determined by subtracting the TSS usage of each bin in WT from the corresponding bin in mutants. The upstream and downstream shifters are colored purple and orange, respectively. **c**. Primer extension and quantification showing analyzed *tfb3* MPA^S^ (Left) and His^+^ (Right) mutant effects on *ADH1* TSS distribution. **d.** Primer extension and quantification showing analyzed *tfa1* MPA^S^ and His^+^ mutant effects on *ADH1* TSS distribution. Color coding of the *tfb3* and *tfa1* allele class in this graph is used throughout the figures. Shades of purple represent upstream shifting *tfa1* or *tfb3* alleles when used to annotate figures. Shades of orange represent downstream shifting *tfa1* or *tfb3* alleles. Averages of ≥3 biological replicates ± standard deviation are shown.

### *tfb3* alleles show mostly additive interactions with *ssl2* mutants and *sub1*Δ

To investigate the relationships between putative *ssl2* processivity mutants and *tfb3* alleles, we generated *tfb3/ssl2* double mutants that combined both upstream- and downstream-shifting variants of the two genes. When combining upstream-shifting *tfb3* alleles with upstream-shifting *ssl2* alleles we observed extensive synthetic sickness or lethality, consistent with each allele conferring defects independently and the double mutants having stronger phenotypes (**Figures 3a, S2a**). When combining *tfb3* and *ssl2* alleles of opposing phenotypes and TSS shifts, we observe that there is partial mutual suppression between the upstream *tfb3-K17E* allele and downstream shifting *ssl2* alleles. The *tfb3-K17E* allele shows moderate to strong suppression of the *ssl2* allele His^+^ phenotype but there is weaker suppression of the *tfb3-K17E* MPA^S^ phenotype in the doubles. This result is consistent with some additivity and that *tfb3-K17E* is either a stronger upstream-shifting allele than the *ssl2* alleles are downstream, or there is partial epistasis of *tfb3-K17E* to the tested *ssl2* alleles (*ssl2*-*N230I* and *ssl2-R636C*) (**Figure 3b top**). Additionally, in a patch phenotyping assay for double mutants combining *ssl2* His⁺ alleles (N230I and R636C) with *tfb3* alleles exhibiting varying strengths of MPA^S^ phenotype, we observed a spectrum of suppression for transcriptional phenotypes. Specifically, double mutants carrying the strong *tfb3* MPA^S^ alleles (C16R, S23P, L22P, and C39R) showed suppression of both the MPA^S^ phenotype of the *tfb3* single mutants and the His⁺ phenotype of the *ssl2* single mutants **(Figure S2a**).

**Figure 3:**
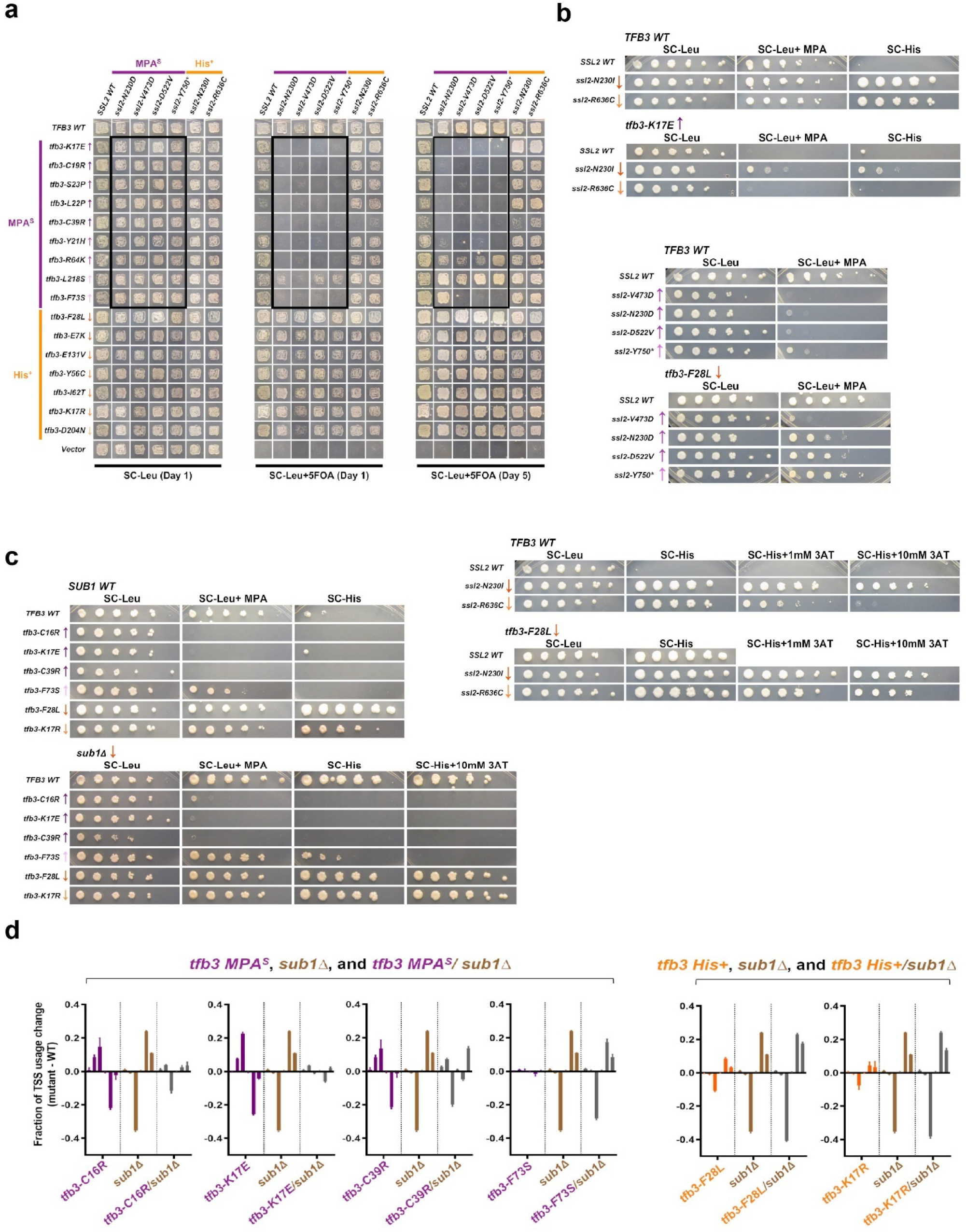
Genetic interactions of *tfb3* alleles with *ssl2* alleles or the PIC cofactor *sub1*Δ show additive interactions. **a.** Patch assay showing viability of *tfb3/ssl2* double mutants assessed by plasmid shuffling. The left panel shows growth on SC-Leu plate which is the control state where WT *TFB3* is present. Middle and right panels show growth on SC-Leu+5FOA plate on day 1 and day 5. Upward and downward arrows denote previously detected or predicted single-mutant effects on TSS distribution (up arrow indicating upstream shifts, down arrow indicating downstream shifts), with greater color intensity indicating stronger phenotypic effects. Patches in black boxes are double mutants that combine *tfb3* MPA^S^ mutants and *ssl2* MPA^S^ mutants**. b.** Spot assay results for double mutants combining *tfb3 K17E* (MPA^S^ allele) and *ssl2* His^+^ alleles (Top), *tfb3 F28L* (His^+^ allele) and *ssl2* MPA^S^ alleles (middle) and *tfb3 F28L* (His^+^ allele) and *ssl2* His^+^ alleles (bottom). **c.** Spot assay results for double mutants combining *tfb3* alleles (both MPA^S^ and His^+^) and *sub1*Δ. **d.** Quantification of primer extension result showing TSS usage changes of single and double mutants from WT. Signals for *tfb3* MPA^S^, *tfb3* His^+^, *sub1*Δ, and *tfb3/sub1*Δ mutants are shown in purple, orange, brown, and gray, respectively. Averages of ≥3 biological replicates ± standard deviation are shown.

When combining upstream shifting *ssl2* alleles with downstream shifting *tfb3-F28L* we observed moderate suppression of the *ssl2* alleles’ MPA^S^, consistent with additivity between these alleles. This additive interaction was further supported by patch phenotyping of double mutants combining *ssl2* MPA^S^ alleles with *tfb3* His^+^ alleles exhibiting different strengths of the corresponding phenotypes. Specifically, double mutants carrying strong or moderate *tfb3* His⁺ alleles together with *ssl2* MPA^S^ alleles displayed weaker MPA^S^ phenotypes compared to the corresponding *ssl2* single mutants (**Figures 3b, S2b**). Finally, we also observe stronger His^+^ phenotypes when His^+^ alleles of *ssl2* are combined with *tfb3-F28L*, also consistent with additivity (**Figure 3b bottom**). Similarly, additive interaction of *ssl2* His^+^ with *tfb3* His^+^ was further supported by patch phenotyping where combination of the *ssl2*-*R636C* with other *tfb3* His^+^ alleles showed stronger His^+^ phenotypes than *ssl2-R636C* single (**Figure S2c**). These double mutant analysis results are in contrast with Pol II allele genetic interactions with *ssl2* alleles, where epistasis was much more prominent and there was very little synthetic lethality (ZHAO *et al*. 2021). In other words, we interpret effects of *tfb3* alleles as potentially altering scanning processivity independently or in parallel with *ssl2* allele effects on processivity, but we can’t rule out additional effects on initiation efficiency or other inputs to scanning (See Discussion).

We also asked about genetic interactions between *sub1*Δ and *tfb3* alleles. Sub1, the yeast homolog of PC4 (HENRY *et al*. 1996; KNAUS *et al*. 1996), is a PIC-associated protein (TIMOTHY *et al*. 2011) that can shows extensive genetic interactions with PIC components (KNAUS *et al*. 1996; WU *et al*. 1999; CALVO AND MANLEY 2005; JIN AND KAPLAN 2015; GARAVÍS *et al*. 2017; ZHAO *et al*. 2021). Deletion of *SUB1* (*sub1*Δ) causes a significant downstream shift in TSS distributions in yeast at *ADH1* (KOYAMA *et al*. 2008; BRABERG *et al*. 2013). Moreover, its distinct genetic interactions with Pol II or *sua7-1* alleles compared with *ssl2* alleles suggested that Sub1 might function in controlling scanning processivity, with loss of Sub1 leading to an increase in processivity (BRABERG *et al*. 2013; JIN AND KAPLAN 2015; ZHAO *et al*. 2021). When *sub1*Δ is combined with upstream-shifting *tfb3* alleles, *tfb3* alleles’ MPA^S^ phenotypes appeared to be mostly unaffected and the *sub1*Δ His^+^ phenotype was suppressed. These phenotypes report on *IMD2* TSS usage and interpretation from these phenotypes alone would suggest *tfb3* alleles were epistatic to *sub1*Δ (**Figure 3c**). However, examination of primer extension for *ADH1* TSS distributions suggests that in fact *sub1*Δ and *tfb3* upstream-shifting alleles can be additive. When combining *sub1*Δ with downstream shifting *tfb3* alleles, His^+^ plate phenotypes of double mutants were difficult to differentiate from *sub1*Δ alone, because it’s His^+^ phenotype is already so strong. However, examination of *ADH1* TSS distributions in double mutants shows that double mutants have slightly greater effects than *sub1*Δ alone, consistent with additivity between *sub1*Δ and *tfb3* downstream-shifting alleles (**Figures 3d, S2d**).

### *rpb7* and *tfb3* alleles at the Tfb3-Rpb7 interface show strong initiation phenotypes

To further study the mechanism of Tfb3 function in TSS selection, we focused our attention on the Tfb3-Rpb7 interface. The Cryo-EM structure of the budding yeast PIC from Schilbach *et al*. 2021 revealed the TFIIE-TFIIH-Pol II interactions within the PIC (**Figures 1e**, **4a**) (SCHILBACH *et al*. 2021). Our genetic screens suggest that Tfb3-Tfa1 interfaces play a critical role in TSS selection during promoter scanning, likely by contributing to scanning processivity. In addition, structural studies and limited mutational analysis previously pointed to potential salt bridges between Tfb3 and Rpb7 as being critical for normal initiation by promoter scanning (BRABERG *et al*. 2013; SCHILBACH *et al*. 2017; SCHILBACH *et al*. 2021; YANG *et al*. 2022). A salt bridge connects Tfb3 R64 to Rpb7 D166 (which we will refer to as salt bridge 1 (SB-1), and a second salt bridge connects Tfb3 K65 to Rpb7 E165 (SB-2) (SCHILBACH *et al*. 2017) (**Figure 4b, left**). However, in a more recent cryo-EM structure only SB-1 between Tfb3 R64 and Rpb7 D166 was observed (**Figure 4b, right**) (SCHILBACH *et al*. 2021). To further dissect this interface, we performed systematic genetic analyses to determine whether one or both salt bridges functionally contribute to scanning.

**Figure 4:**
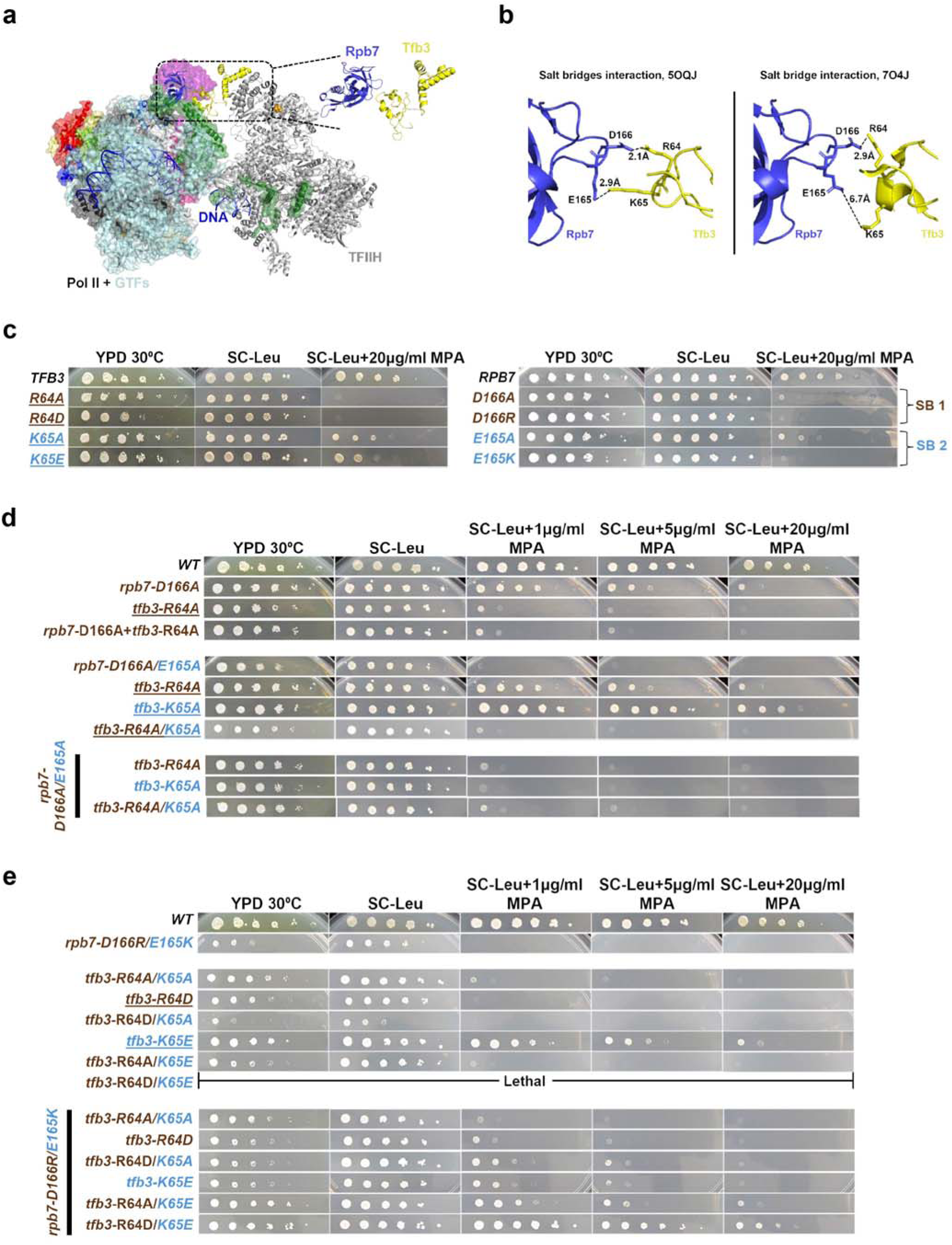
Genetic dissection of Rpb7-Tfb3-Tfa1 interface. **a**. Position of Rpb7-Tfb3-Tfa1 interface to TFIIH and Pol II in the yeast PIC (PDB 7O4J)(SCHILBACH *et al*. 2021). **b**. Interaction between Rpb7 and Tfb3 via two putative salt bridges (left, PDB 5OQJ) (SCHILBACH *et al*. 2017) vs single putative salt bridge (right, PDB 7O4J) (SCHILBACH *et al*. 2021) in distinct cryo-EM structures of yeast PIC. Rpb7 and Tfb3 are colored yellow and blue, respectively. Residue numbers are color-coded according to their parent protein color. **c.** Mutants at Rpb7-Tfb3 interacting surfaces show strong MPA^S^ phenotypes. Spot assay results for *tfb3* (left) and *rpb7* (right) alleles on YPD, SC-Leu and SC-Leu+20 μg/ml MPA. WT is labeled in black, mutants present at putative SB-1 are labeled in brown, mutants at putative SB-2 are labeled in orange, with previously described mutants underlined. **d.** Double mutant combination of alanine-substituted *rpb7* and *tfb3* alleles at either putative salt bridges or both, are epistatic. **e**. Charge reversal at both putative salt bridges at the Tfb3-Rpb7 interface suppresses individual mutant initiation defects. Lower MPA concentrations (1 and 5 μg/ml) were used in panels **e–f** to reveal subtle differences in phenotype strength that are masked at 20 μg/ml MPA.

To determine the contributions for putative salt bridge forming residues at the Rpb7-Tfb3 interface to TSS selection, we generated a battery of *rpb7* and *tfb3* single mutants and *rpb7/tfb3* double mutants (Materials and Method). To enable ease of double mutant generation, we employed a plasmid-shuffle approach where *tfb3* mutants were generated using site-directed mutagenesis in *TFB3 LEU2* plasmids and *rpb7* mutants were generated in the genome by CRISPR. *TFB3* or *tfb3* plasmids were introduced into relevant *RPB7* or *rpb7* strains deleted for *TFB3* by transformation, followed by selection against a WT *TFB3 URA3* plasmid (**Figure S3a**).

Alleles with substitutions at either one or both salt bridge-forming residues at this interface were screened for initiation phenotypes using those employed for genetic screening. We found that mutations in all individual residues within the putative Rpb7-Tfb3 salt bridges conferred MPA sensitivity, which is predictive of upstream TSS shifts relative to WT. Mutations in SB-1 residues were generally more MPA sensitive and exhibited stronger overall growth defects than those in SB-2. Similarly, for both *tfb3* and *rpb7* mutants, charge-altering substitutions caused more severe phenotypes (both in growth and MPA sensitivity) than alanine substitutions (**Figure 4c**). We interpret this as consistent with charge incompatibility arising when only one partner of the putative salt bridge is altered, resulting in the introduction of like charges in proximity. We next sought to test this model more rigorously through an extensive series of double-mutant analyses.

### Genetic dissection of mutants at the Tfb3-Rpb7 interface supports functional interactions between Tfb3-Rpb7 via two pairs of amino residues observed as salt bridges in different cryo-EM structures

To determine if Tfb3 and Rpb7 residues work together through physical interactions to facilitate promoter scanning in *S. cerevisiae*, we evaluated the genetic interactions of *tfb3/rpb7* double mutants and the potential restoration of growth when one or both salt bridges were restored but in novel configurations through charge reversal mutations. First, to assess the nature of the genetic interactions—additive, suppressive, or epistatic—at the predicted salt bridge–forming sites, we performed phenotypic analyses of double mutants harboring alanine-substituted *rpb7* alleles in combination with alanine-substituted *tfb3* alleles (MANI *et al*. 2008). We observed that combining either single or double alanine-substituted *rpb7* alleles at putative SB-1 or SB-2 or both with either single or double alanine-substituted *tfb3* alleles show epistatic interactions (**Figure 4d**). These results are consistent with these residues requiring each other for function, and mutating one side of the interaction means that mutation of the other side does not confer a further defect.

Next, to determine whether the Rpb7-Tfb3 interface relies on one or both salt bridges, we performed charge identity reversal experiments at SB-1, SB-2, or both simultaneously. We hypothesized that only complete charge reversal at both putative salt bridges would restore the transcriptional defects caused by mutations that compromise both sites. As a starting point, we characterized the growth and MPA phenotypes of the individual *rpb7* and *tfb3* alleles used to construct the combined mutants (**Figure 4e, top**). The *rpb7-D166R/E165K* mutant, in which both salt bridge–forming residues (D166 and E165) are converted to positively charged residues, is in direct opposition to the positively charged partners of Tfb3 (R64 and K65). This mutant exhibits a strong MPA^S^ phenotype and a pronounced growth defect. We therefore used *rpb7-D166R/E165K* as the reference “sick” allele in plasmid shuffle experiments designed to evaluate whether restoring charge complementarity at the Tfb3 positions could suppress its MPA^S^ and growth phenotype.

The charge reversal at a single putative salt bridge provided only limited suppression of the initiation defect of the *rpb7-D166R/E165K* mutant. For example, the combinations *rpb7-D166R/E165K* with *tfb3-R64D* (putatively restoring SB-1) and *rpb7-D166R/E165K* with tfb*3-K65E* (putatively restoring SB-2) yielded only modest suppression in growth and MPA sensitivity compared to *rpb7-D166R/E165K* alone. To account for potential charge incompatibility from having two adjacent positively charged residues, we generated quadruple mutants designed for charge reversal of a single salt bridge but with an alanine substitution in one of the pair of residues supporting the other salt bridge: *rpb7-D166R/E165K* + *tfb3-R64D/K65A* (restoring SB-1 but removing charge incompatibility at SB-2) and *rpb7-D166R/E165K* + *tfb3-R64A/K65E* (restoring SB-2 but removing charge incompatibility at SB-1). These mutant combinations displayed improved growth phenotypes relative to their charge-incompatible counterparts were still not wild type (**Figure 4e, bottom**). Additionally, to remove charge or side chain effects from each of the salt bridge pairs and examine the relative contributions of SB-1 and SB-2 more carefully, we constructed charge reversal mutations for each salt bridge while alanine substituting both residues of the other salt bridge. These combinations included *rpb7-D166R/E165A* or *rpb7-D166A/E165K*, which were combined with various *tfb3* alleles. Quadruple mutants such as r*pb7-D166R/E165A + tfb3-R64D/K65A* (restoring SB-1 only) and *rpb7-D166A/E165K + tfb3-R64A/K65E* (restoring SB-2 only) showed partial suppression of the double-salt-bridge defect (**Figure S3b**), consistent with the results obtained from *rpb7-D166R/E165K + tfb3-R64D/K65A and rpb7-D166R/E165K + tfb3-R64A/K65E*. Finally, the complete charge reversal at both salt bridges achieved by combining *rpb7-D166R/E165K* with *tfb3-R64D/K65E*, suppressed the severe growth defect and suppressed the MPA^S^ phenotype of the *rpb7-D166R/E165K* mutant (**Figure 4e**). In this case, restoration of charge complementarity at both interfaces was sufficient to suppress the most growth defects.

To further confirm the charge-based nature of the Tfb3-Rpb7 interaction, we generated an alternative mutant, *rpb7-D166K/E165R*, which reverses the charge pattern with different substitutions. When combined with the corresponding charge-reversal *tfb3* alleles, the combined mutant exhibited suppression comparable to *rpb7-D166R/E165K*, reinforcing the functional relevance of charge complementarity at this interface (**Figure S3c**). Collectively, these genetic data indicate that both salt bridges contribute to a stable and functional Tfb3-Rpb7 interaction, and that disruption of either alone is sufficient to perturb normal initiation.

### *rpb7* alleles at Tfb3-Rpb7 salt bridge forming interface show genetic interactions with additional *tfb3* alleles

To further investigate the nature of the Tfb3-Rpb7 interaction, we examined the genetic interactions of *rpb7* alleles at interface with novel *tfb3* alleles outside the Tfb3-Rpb7 interface. The strong His^+^ *tfb3-M12K* shows slight suppression of MPA^S^ alleles at Rpb7-Tfb3 interface. The *tfb3-M12K* suppresses the growth defect and MPA^S^ phenotype of *rpb7-D166A* whereas it slightly suppresses growth defect of *rpb7-D166A/E165A* (**Figure 5**). This suppression of MPA^S^ interface mutants by His^+^ mutants support the scanning defect at Rpb7-Tfb3 is plastic in nature and that His^+^ mutants may indeed be acting by increasing interactions. The MPA^S^ mutants disrupting salt bridge interactions at Tfb3-Rpb7 interface such as *rpb7-D166R/E165K*, *rpb7-D166A/E165A* and *tfb3-R64A/K65A* are strong MPA^S^ and had growth defects (**Figure 4**) but they are still viable. This observation suggests that the defect at Tfb3-Rpb7 interface still supports some scanning in vivo. However, when we combine the *rpb7* alleles at Tfb3-Rpb7 interface with *tfb3* MPA^S^ alleles elsewhere, we observed additive effects. When the sick MPA^S^ *tfb3-C39R* allele was combined with *rpb7-D166A* or *rpb7-D166A/E165A,* we observed lethality for both combinations. Additionally, with *tfb3-Y21H*, when combined with *rpb7-D166A*, we observed strong additive effects in MPA^S^ phenotype and growth defects. When we combined *tfb3-Y21H* with *rpb7-D166A/E165A,* lethality was observed, consistent with an additive interaction (**Figure 5**).

**Figure 5:**
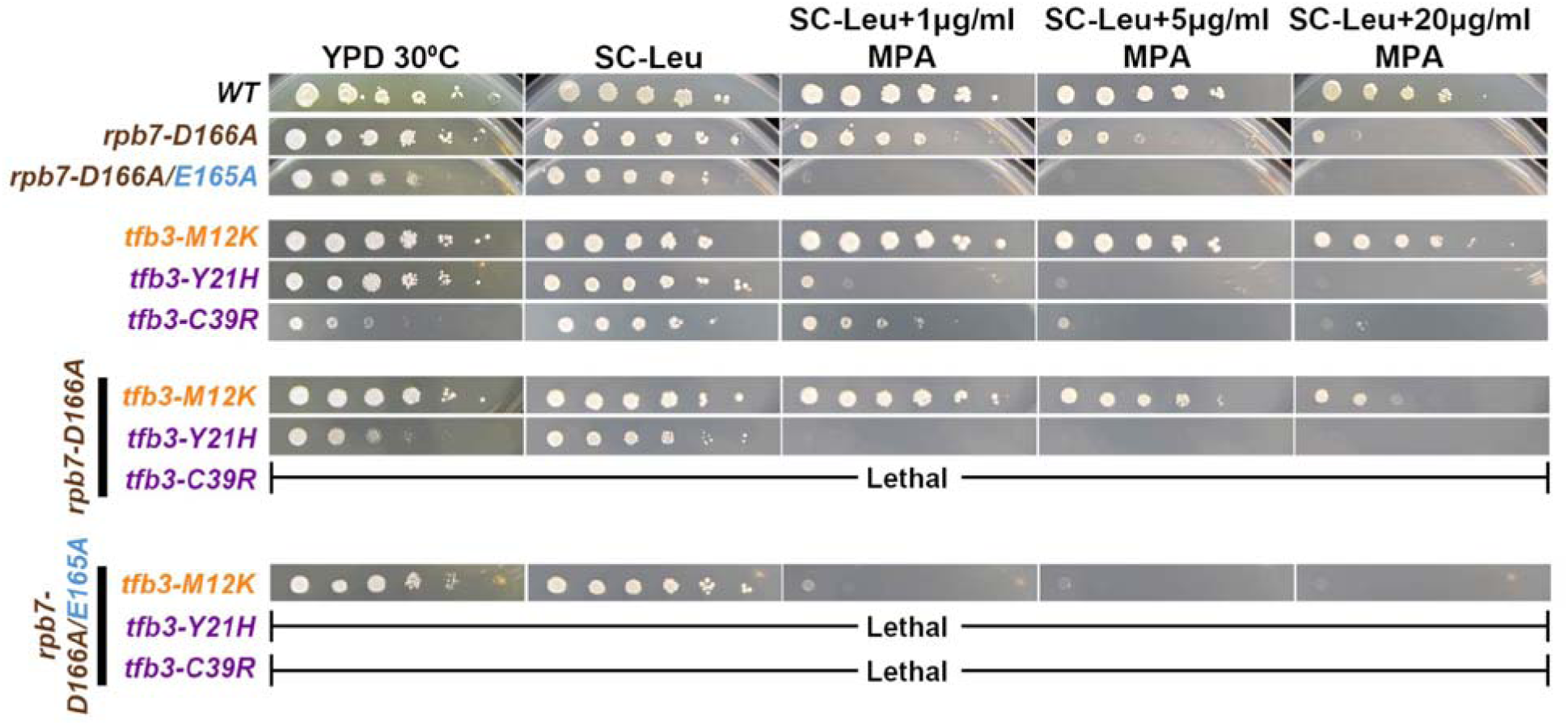
*rpb7* mutants in a salt bridge-forming residue of Rpb7-Tfb3 show additive interactions with upstream and downstream shifting *tfb3* alleles outside the Rpb7-Tfb3 interface. Spot assay results for double mutant combination of *rpb7-D166A* or *rpb7-D166A/E165A* with upstream shifting *tfb3* alleles (purple) or a downstream shifting *tfb3* allele (orange).

### Genome-wide sequencing (TSS-seq and STRIPE-seq) identifies global *tfa1*, *tfb3*, and *rpb7* initiation effects in *S. cerevisiae*

We examined the 5′ ends of RNA transcripts for select *tfb3*, *tfa1* and *rpb7* alleles on a genome wide level. For a subset of representative mutants isolated through genetic screens we performed TSS-Seq (VVEDENSKAYA *et al*. 2015), and a subset of those mutants were additionally validated by STRIPE-Seq (POLICASTRO *et al*. 2020) (a CAGE-related protocol for TSS mapping) in parallel (**Figures S4a**, **S4b**, **Materials and Methods**). We also analyzed salt bridge alleles by STRIPE-Seq (see below). In total, six *tfb3* alleles, three MPA^S^ (K17E, C39R, and R64K) and three His^+^ (K17R, F28L, and I62A)), four *tfa1* alleles, two MPA^S^ (K187E, and I183T) and two His^+^ (F14L, and K99I) and one *rpb7* (MPA^S^, D166G) allele were analyzed along with their strain-specific WT controls by TSS-Seq. The STRIPE-Seq analyzed strains included two *tfb3* (MPA^S^; R64K and His^+^; F28L) and two *tfa1* (MPA^S^; K187E and His^+^; K99I) along with their strain-specific WT controls. The positions and counts of the 5′ ends of uniquely mapped reads were extracted to assess reproducibility between biological replicates. We observed strong correlations among replicates for both TSS-Seq and STRIPE-Seq datasets (**Figures 6a**, **S4c, S5a, S5b**). As described in Qiu et al. 2020 and Zhao et al. 2021, we investigated 5979 promoters across the genome and after stringent data filtering and read depth thresholding (**Materials and Methods**), we retained 3226 promoters for TSS-Seq and 2883 for STRIPE-Seq (QIU *et al*. 2020; ZHAO *et al*. 2021).

**Figure 6:**
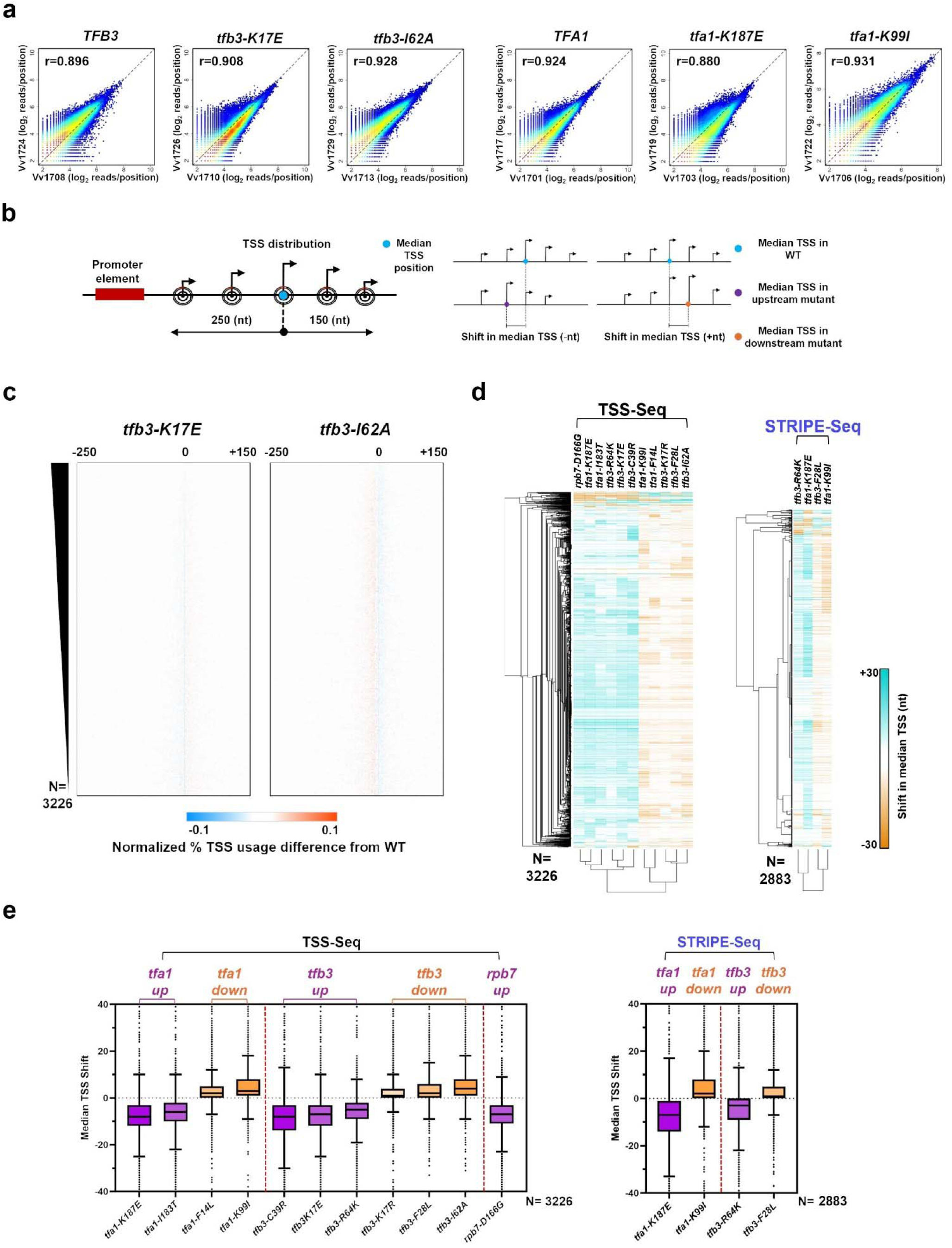
The *tfb3*, *tfa1*, and *rpb7* allele shift transcription start site (TSS) distributions genome wide. **a.** Scatter plot showing the correlation of log_2_ transformed reads at individual genome positions for all positions ≥ 3 reads for biological replicates shown for representative alleles. **b.** Schematic depicting the TSS distribution within a promoter window centered on the median of the wild-type (WT) TSS distribution (Left). Schematic showing shift of the median TSS, either upstream or downstream, in a mutant relative to the WT (Right). **c.** Heatmaps of TSS-normalized read differences between WT and two *tfb3* (*tfb3-K17E*; upstream shifting and *tfb3-I62A*; downstream shifting) mutants within defined promoter windows. Promoter classes were rank ordered according to total reads in WT. The promoter-normalized read differences between the mutant and WT are represented using a color gradient from cyan (negative) to orange (positive). **d.** Heatmap with hierarchical clustering of median TSS shifts in *tfa1*, *tfb3*, or *rpb7* mutants generated by TSS-Seq (Left) and a subset of mutants examined by STRIPE-Seq (Right). **e.** Box plots showing median TSS shifts across promoters in *tfa1*, *tfb3*, or *rpb7* mutants, TSS-Seq (Left) and a subset of mutants with STRIPE-Seq (Right). Median TSS shifts across promoters, are statistically distinguished from wildtype at p < 0.0001 for all mutants (Wilcoxon signed rank test).

To examine how these *tfb3*, *tfa1*, and *rpb7* alleles affect transcription start site (TSS) distributions, we applied two main metrics for TSS usage as described in Qiu et al. 2020 and Zhao et al. 2021 (QIU *et al*. 2020; ZHAO *et al*. 2021). First, we quantified TSS distribution differences between WT and mutants within defined 401 bp promoter windows (**Figure 6b, left**). We observed that all MPA^S^ alleles, such as *tfb3-R64K*, *tfa1-K187E*, and *rpb7-D166G*, exhibited upstream shifts in the median position of the TSS distribution on average across analyzed promoters. In contrast, His⁺ alleles for both *tfa1* and *tfb3* displayed increased TSS usage at downstream sites, consistent with downstream shifts in initiation (**Figures 6b**, **6c**). Next, to quantify changes in TSS usage, we used a metric defined as the “median TSS shift”, calculated by subtracting the WT median TSS position within each promoter window from that of the mutant (**Figure 6b, right**). For each mutant, we show the median TSS shifts across promoters (**Figures 6d**, **6e**). Consistent with genetic and primer extension observations; *tfb3*, *tfa1* and *rpb7* MPA^S^ alleles shift the median TSS positions upstream whereas His^+^ alleles shift median TSS positions downstream at the majority of promoters (**Figures 6d**, **6e, S5a)**. As both MPA^S^ and His^+^ alleles show a gradient of phenotypic strength in genetic tests, the same holds true for median TSS shift. For example, Stronger MPA^S^ mutants like *tfb3-C39R* and *tfa1-K187E* shows stronger median TSS shift in upstream direction, with the same pattern holding true for His^+^ alleles (**Figures 1c**, **1d**, **6e**). These results are consistent with phenotypes of mutants being predictive of global TSS defects. TSS-Seq and STRIPE-Seq techniques both recapitulate the observed effect (**Figures 6d**, **6e**). Analysis of individual promoters for statistically significant shifts relative to WT indicates a large directional bias in number of promoters significantly shifted for each mutant, reflective of the overall biases in shifts observed in aggregate (**Figure S6**).

### Genome-wide analysis of TSS selection via STRIPE-seq corroborates interaction between Tfb3-Rpb7 via two salt bridges

The observed growth suppression of the sick *rpb7-D166R/E165K* allele by charge reversal *tfb3-R64D/K65E* suggests potential widespread effects on the *rpb7-D166R/E165K* phenotype. However, the MPA phenotype assessed derives from defects at a single promoter, *IMD2*. To assess potential suppression of defects genome wide, we analyzed a series of mutants by STRIPE-seq (**Figure 7**). Tested alleles included *tfb3-R64K* (SB-1 impaired), *tfb3-R64A/K65A* (both salt bridges impaired from the Tfb3 side), *rpb7-D166A/K165A* (both salt bridges impaired from Rpb7 side), *rpb7-D166R/E165K* (a slow growth allele with both salt bridges impaired along with charge incompatibility) and *rpb7-D166R/E165K+tfb3-R64D/K65E* (quadruple mutant with charge reversal). As expected, these alleles shift the median TSS positions upstream across the majority of promoters, consistent with their MPA^S^ phenotypic strength (**Figures 7a**, **7b**). The alleles where both salt bridge interactions were impaired showed strong shift in median TSS upstream whereas the quadruple mutant with the potential for salt bridge restoration was much closer to wild type with a median shift near zero. This observation validates our salt bridge interaction, where charge reversal allele strongly suppresses the upstream shift in TSSs for the sick *rpb7-D166R/E165K* allele at the majority of promoters in genome wide level.

**Figure 7:**
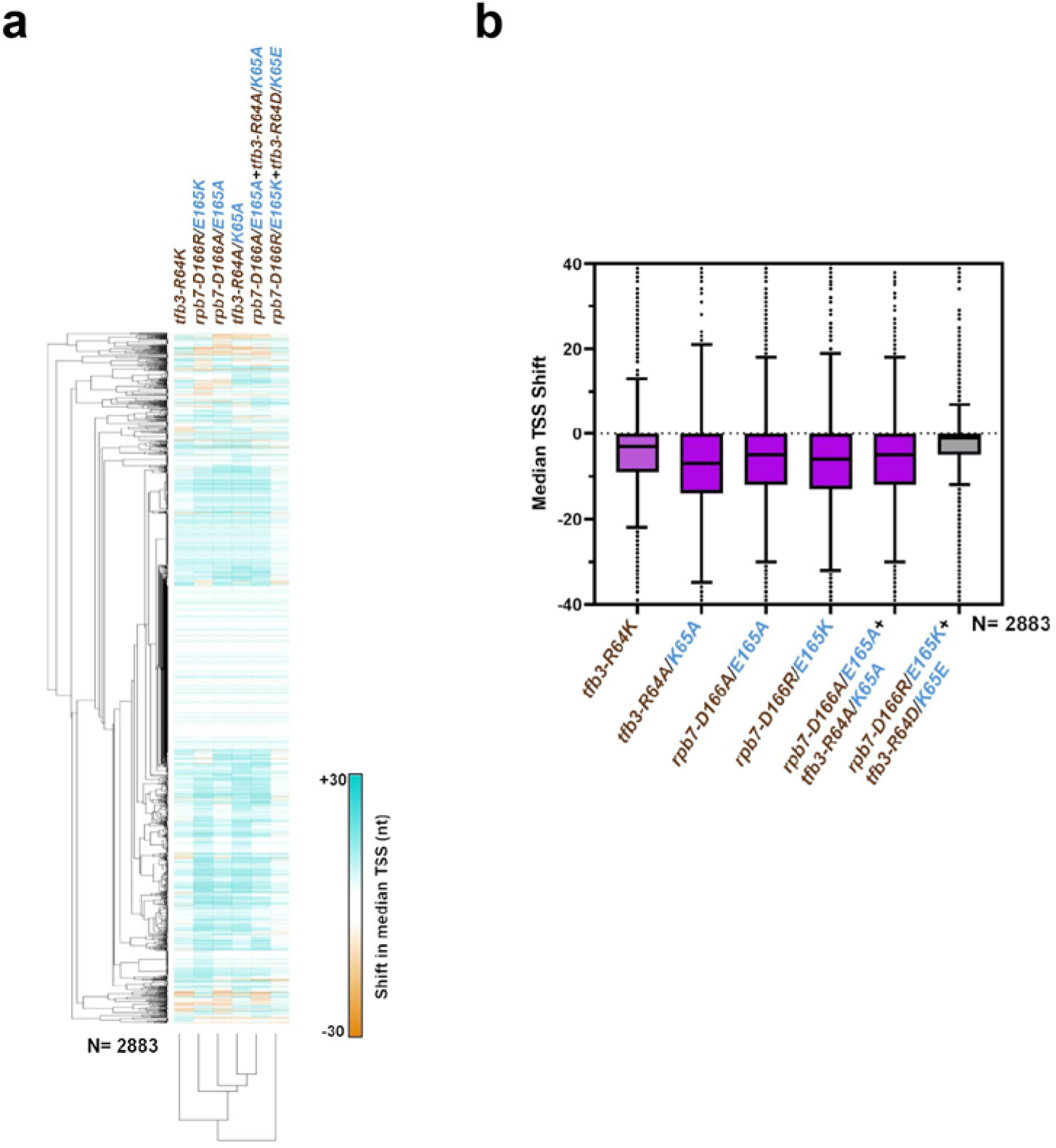
STRIPE-seq validates salt bridge interaction as charge reversal partially restores upstream shift of TSSs in sick *rpb7* mutant. **a.** Heatmap with hierarchical clustering of median TSS shifts in *tfb3*, or *rpb7* salt bridge mutants. **b.** Box plot showing median TSS shifts across promoters in *tfb3*, or *rpb7* salt bridge mutants. Charge reversal mutant (grey) suppresses the median TSS shift. Median TSS shifts across promoters, are statistically distinguished from wildtype at p < 0.0001 for all mutants (Wilcoxon signed rank test).

### Qualitative similarity between TSS sequencing techniques but technique-specific differences

Our experiments assessed two types of TSS mapping techniques. TSS-seq as employed by our group, starts with rRNA-depleted RNA and uses a relatively large amount of input (∼1 ug), but is highly reproducible and generates deep promoter coverage (CHO *et al*. 2009; MITSCHKE *et al*. 2011; VVEDENSKAYA *et al*. 2015; QIU *et al*. 2020; ZHAO *et al*. 2021). STRIPE-seq is a CAGE-related protocol that utilizes template switching for addition of a 5′ adapter and second strand synthesis (POLICASTRO *et al*. 2020). An advantage of STRIPE-seq is that it calls for a much smaller amount of starting material – as low as 50 ng total RNA. However, for our purposes, we started with ∼150-200 ng of rRNA-depleted RNA, which is still 5-fold less than for our TSS-seq. STRIPE-seq showed strong agreement with TSS-seq in capturing the expected direction of TSS shifts for tested alleles, although biological replicates showed some variability in median shift (**Figure S5a, left**), and fewer promoters were above threshold for consideration compared to TSS-seq. To directly compare the two methods, we analyzed 2,564 promoters detected by both. Within this shared set, mutants exhibited consistent directional TSS shifts, but the magnitude of shifts differed between techniques (**Figures S7b, S7d**). Clustering analysis of the median TSS positions revealed that data points derived from each technique grouped separately (**Figure S7a**). Similarly, for median TSS shift, although upstream- and downstream-shifting alleles formed distinct clusters, the separation appeared to be driven primarily by the sequencing technique rather than by allele-specific effects (**Figures S7b**). Principal component analysis of the median TSS shifts further confirmed that variation was largely technique-dependent (**Figure S7c**). Together, these results demonstrate that STRIPE-seq reliably captures qualitative TSS-shift behavior of mutants, but differences due to each technique suggest they can be compared qualitatively and not quantitatively.

## DISCUSSION

Our genetic and genomic results reveal a notable plasticity in how tested GTFs can contribute to promoter scanning. For both TFIIE subunit Tfa1 and the architectural TFIIH component Tfb3, we were able to obtain mutants that altered TSS distributions in either the upstream or downstream direction.

For previously examined Pol II and *SSL2* alleles, the enzymatic activity of each provided a straightforward explanation for finding alleles of different classes – that they increased or decreased activity relative to WT. Here, our studies show that additional GTFs may also increase or decrease the outcomes of scanning as evidenced by shifts in TSS distributions. Consistent with our targeted genetic screens revealing mutants in GTF components able to alter TSS distributions, our other work using unbiased genetic selections across the genome have also revealed mutants across GTFs and in multiple components of TFIIH able to shift TSSs (ARORA *et al*. 2025).

Previous genetic studies identified Rpb7 and Tfb3 alleles at the Tfb3-Rpb7 interface that conferred MPA^S^ (BRABERG *et al*. 2013; YANG *et al*. 2022; YANG *et al*. 2024). The location and nature of those alleles suggest that defects in protein interactions confer the MPA^S^ phenotype, and we interpret this phenotype as, minimally, a reduction in TFIIH scanning processivity leading to inability for scanning to reach downstream TSSs. These results would be consistent with this conserved interface being important for the coupling of TFIIH translocation to continued TFIIH scanning in yeast. However, our genetic screens have isolated His^+^ alleles able to shift TSS usage downstream for both Tfa1 and Tfb3. As noted above, Arora et al. 2025 also identified His^+^ alleles for the TFIIH components Rad3, Tfb2, Tfb4 and Ssl1 (ARORA *et al*. 2025).

How do these novel alleles alter scanning? Given that we interpret the converse phenotype (MPA^S^) for alleles of these genes as compromising the physical integrity of the Tfa1-Tfb3-Rpb7 interface, we interpret His^+^ alleles as strengthening the interface or adding plasticity to it. This interpretation suggests roles for the architectural components of TFIIH in the PIC that are not enzymatic components themselves but can feedback or modulate the biochemical activities involved in promoter scanning, which are Pol II catalysis and TFIIH translocation. Our genetic interactions between interface components and *ssl2* alleles or *sub1*Δ suggest that *tfb3* alleles at least are additive with *ssl2* and *sub1*Δ, suggesting that some of their effects alter TFIIH processivity. However, upstream shifting *rpb7, tfa1,* and *tfb3* alleles tested at *ADH1* all appear stronger than tested *ssl2* alleles (**Figure 2**), though weaker than strong upstream shifting Pol II alleles (KAPLAN *et al*. 2012; BRABERG *et al*. 2013; QIU *et al*. 2016; ZHAO *et al*. 2021). A simple explanation is that these observations reflect quantitative differences in allele strength. This explanation is somewhat unappealing because an allele that only affects scanning processivity would truncate downstream scanning, therefore leading to relative increases in upstream TSSs. Processivity mutants would not be predicted to alter actual initiation efficiencies at any position within the scanning window, just the length of the scanning window. In contrast, Pol II and TFIIF alleles that shift TSSs upstream appear to strongly activate the use of upstream TSSs and this is consistent with their ability to alter efficiency of each TSS within the scanning window. An intriguing alternative is that Pol II-TFIIH interface mutants might communicate with the Pol II active site (affecting innate initiation efficiency), the TFIIH active site (affecting scanning rate), or an additional PIC activity during scanning. Such an additional activity could reflect the unknown mechanism that restricts initiation close to the core promoter/TATA element. Relief of this restriction would also be predicted to activate TSS usage at upstream positions without needing to directly communicate to the Pol II active site.

Together, these findings suggest that multiple components of the PIC, both enzymatic and structural, participate in shaping the dynamics of promoter scanning. The ability of non-enzymatic factors and interfaces to influence scanning underscores the plasticity of the scanning process. The fact that scanning is easily affected in either direction by individual mutations is consistent with the types of changes that may have contributed to the evolution of promoter scanning within otherwise highly conserved and essential components of the eukaryotic PIC.

## MATERIALS AND METHODS

### Yeast strains and media

Yeast strains are derived from a GAL^+^ of S288C (FY2) (WINSTON *et al*. 1995). Yeast strains used in this study for *the tfb3* mutant screen, *tfb3/ssl2* double mutants, *tfb3/sub1*Δ *double mutants*, *tfa1* mutant screen, *rpb7* and *tfb3* mutants at the Tfb3-Rpb7 interacting surface, and related experiments are listed in **Supplementary Table 1.** Yeast media are prepared following standard protocols (AMBERG 2005). YP solid medium is made of yeast extract (1% w/v; BD), peptone (2% w/v; BD, 211677) and Bacto agar (2% w/v; BD, 214010), supplemented with adenine (0.15 mM; Sigma-Aldrich, A9126) and L-tryptophan (0.4 mM; Sigma-Aldrich T0254). YPD plates contained dextrose (2% w/v; VWR, VWRBK876), YPRaf plates contained raffinose (2% w/v; Amresco, J392), and YPRaf/Gal plates contained raffinose (2% w/v; Amresco, J392) and galactose (1% w/v; Amresco, 0637) as carbon sources. YPRaf and YPRaf/Gal plates also contain antimycin A (1 μg/ml; Sigma, A8674-100mg). Minimal media plates are synthetic complete (“SC”) with amino-acids dropped out as appropriate as described in (AMBERG 2005) with minor alterations as described in (KAPLAN *et al*. 2012): per standard batch formulation, adenine hemisulfate (Sigma-Aldrich, A9126) was 2 g, L-Leucine (Sigma-Aldrich, L8000) was 4 g, Myo-inositol was 0.1 g, para-aminobenzoic acid (PABA) was 0.2 g. SC-Leu+5FOA plates contained 1 mg/ml final concentration of 5-fluoroorotic acid monohydrate (5-FOA, GoldBio, F-230). SC-Leu+MPA plates contained either 20 ug/ml, 5 ug/ml, or 1ug/ml final concentration of mycophenolic acid (MPA, Sigma, M3536-250MG). SC-His+3AT plates contained different final concentrations of 3-Amino-1,2,4-triazole (3-AT, Sigma-Aldrich, A8056).

### Plasmids, bacterial strains, and oligonucleotides

Bacterial strains, plasmids, and oligonucleotides used in this study are listed in **Supplementary Tables 2 and 3**.

### *tfb3* and *tfa1* genetic screening

Novel *tfb3* and *tfa1* mutants were created by PCR-based random mutagenesis coupled with a gap repair. Briefly, to generate randomly mutagenized *tfb3* (*tfb3\**), standard PCR was performed using Taq polymerase (NEB). *tfb3*\* PCR products were then transformed into yeast along with a linearized pRS315-*TFB3* (pCK1664) plasmid with most of the WT TFB3 sequence removed by restriction digest using *Bgl*II and *Nru*I (Thermo Scientific). Leu^+^ transformants were selected. Homologous sequences on each end of the *tfb3*\* PCR products and the gapped *tfb3* vector allowed homologous recombination, resulting in a library of gap-repaired plasmids containing potential *tfb3*\* mutants. Because *TFB3* is an essential gene, these yeast cells are pre-transformed with a pRSII316-*TFB3* (pCK1632) *URA3* plasmid to support growth, while the genomic *TFB3* was deleted to allow plasmid *tfb3* alleles to exhibit phenotypes. After gap repair, cells retaining pCK1632 (pRSII316-*TFB3 URA3*) plasmids were killed when transformants were replica-plated to medium containing 5-FOA (GoldBio). Yeast cells were then replica-plated to various media to screen for mutants with transcription-related or conditional phenotypes. Plasmids from yeast mutants were recovered and transformed into *Escherichia coli* for amplification, followed by sequencing to identify mutations. Additional phenotyping was performed to determine if the mutant phenotype was plasmid-linked or not. All mutants described here were verified by retransformation into a clean genetic background. A similar approach was taken for the *tfa1* genetic screen.

### Site-directed mutagenesis at Tfb3-Rpb7 interface: CRISPR Cas9 to generate *rpb7* alleles

Single or double amino acids substituted mutants were created targeting Rpb7 residues; D166 and E165, using the CRISPR-Cas9 single plasmid pML107 (LAUGHERY *et al*. 2015). This Leu auxotrophic marker-based pML107 (gift from Francois Robert Lab) allows the expression of a guide RNA (gRNA) and Cas9 gene together. First, the forward (5′ portion contains a sequence that allows gRNA to bind to Cas9) and reverse strands of gRNA targeting the region containing the PAM site located near the site of interest for mutation were designed. Similarly, the forward repair oligo containing the mutated PAM site, and the reverse oligo harboring either single or double substituted amino acids for D166 and E165 were designed. After annealing gRNA oligos, they were cloned into pML107 at *Swa*I and *Bcl*I restriction enzyme digested sites. Finally, 400ng of pML107 expressing Cas9 and gRNA insertion was transformed into yeast cells along with double strand repair oligo (up to 1ug) following the yeast high-efficiency transformation protocol described in (GIETZ AND SCHIESTL 2007). Transformants were grown on selective SC-Leu plates. Several colonies were picked and initially selected for initiation phenotypes at SC-Leu+MPA (20ug/ml) plates. Several MPA^S^ and some MPA^R^ (control) colonies were picked. PCR fragments harboring the PAM site with mutation site, and homology sites were sent for sequencing and verification for desired *rpb7* mutations.

### Site-directed mutagenesis at Tfb3-Rpb7 interface: PCR sewing (Overlap PCR) to generate *tfb3* alleles

*tfb3* mutants at the Tfb3-Rpb7 interface were created by overlap PCR. Two PCR products overlapping at the location of a desired mutation introduced using mutant oligos were used in an additional PCR reaction to generate mutant DNA amplicons with otherwise WT *TFB3* sequence for the insertion into the full length *TFB3* plasmid. Desired *tfb3*\* PCR products were ligated into *Bcl*l and *Smi*l (NEB), digested pRS315 *TFB3 LEU2* plasmid. *tfb3*\* harboring pRS315 *tfb3* LEU2* plasmids were verified by sequencing. Finally, 200ng of each pRS315 *tfb3*\* *LEU2* plasmid was transformed into yeast cells containing pRSII316 *TFB3 URA3* plasmid, while the genomic *TFB3* was removed to allow plasmid *TFB3* alleles to exhibit phenotypes. Transformants were grown on selective SC-Leu plates. Six candidate colonies from SC-Leu plates were patched on SC-Leu and then replicated to a medium containing 5-FOA (GoldBio), to kill cells retaining pRSII316 *TFB3 URA3* plasmids. Clones that have lost URA3 containing plasmid and harbored *tfb3*\* were then saved to the lab collection.

### Primer extension assay

Primer extension (PE) assays to detect putative usage of TSSs in *ADH1* promoter, were performed on 30ug total RNA extracted by using a phenol-chloroform method (SCHMITT *et al*. 1990) per reaction as described in (RANISH AND HAHN 1991) with modifications included in (KAPLAN *et al*. 2012). Primer CKO401 complementary to *ADH1* mRNA was end-labeled with 32P γ-ATP (PerkinElmer) and T4 polynucleotide kinase (Thermo Scientific). A reverse transcription reaction was performed by mixing M-MuLV Reverse Transcriptase (NEB, RNase inhibitor (NEB), dNTPs (GE), DTT, and labeled primer with total RNA. After cDNA formation, RNase A was added to remove excess RNA. Then, the product was run into a sequencing gel that contained 8% acrylamide gel (19:1 acrylamide: bisacrylamide) (Bio-Rad), 1× TBE, and 7 M urea. Finally, the gels were visualized by Molecular Imager PharosFX™ Plus System (Bio-Rad) and quantified by Image Lab (5.2).

### TSS-seq library preparation

Yeast cell cultures were grown in liquid YPD media and cells were harvested at the mid-log phase at a density of 1-1.5 × 10^7^ cells/ml. Cell counts per culture were determined by counting cells with a hemocytometer. We performed library construction for TSS-seq essentially as described by (VVEDENSKAYA *et al*. 2015); the steps are briefly described as follows; 100 μg of the extracted total RNA (phenol-chloroform extraction) was treated with 30 U of DNase I (QIAGEN) and purified using RNeasy Mini Kit (QIAGEN). A riboPOOL probe (siTOOLS Biotech) was used to deplete rRNAs from DNase-treated RNAs. rRNA depletion was done following published riboPOOL Protocol_v1-6 from siTOOLs Biotech which was modified by us to deplete 25ug of DNAse-treated RNAs at once. The rRNA-depleted RNA was resuspended in ∼15 μl of nuclease-free water. From rRNA-depleted RNA, to remove RNA transcripts carrying a 5′ monophosphate moiety (5′-P), ∼1 μg of rRNA-depleted RNA were treated with 1 U Terminator 5′-Phosphate-Dependent Exonuclease (Epicentre) in the 1× Buffer A in the presence of 40 U RNaseOUT in a 50 μl reaction at 30°C for 1 hr. Then, RNA was extracted using the phenol-chloroform method. Next, to get rid of 5′-terminal phosphates, RNAs were treated with 1.5 U CIP (NEB) in 1× NEBuffer three in the presence of 40 U RNaseOUT in a 50 μl reaction at 37°C for 30 min. After the reaction was completed, RNAs were again extracted using the phenol-chloroform method. Now to convert enriched 5′-capped RNA transcripts to 5′-monophosphate RNAs, CIP-treated RNAs were mixed with 12.5 U CapClip (Cellscript) and 40 U RNaseOUT in 1× CapClip reaction buffer in a 40 μl total volume and incubated at 37°C for 1 hr. Then, the reaction went through the phenol-chloroform extraction method. To ligate the 5′ adapter, these CapClip-treated RNA products were mixed with 1 μM 5′ adapter oligonucleotide s1086, 1× T4 RNA ligase buffer, 40 U RNaseOUT, 1 mM ATP, 10% PEG 8000, and 10 U T4 RNA ligase 1 in a 30 μl reaction volume. The reaction was incubated at 16°C for 16 hr and was halted by adding 30 μl of 2× RNA loading dye. The mixtures were then separated by using electrophoresis on 10% 7 M urea slab gels in 1× TBE buffer. Then the gel was incubated with SYBR Gold nucleic acid gel stain. RNA products migrating above the 5′ adapter oligo were recovered from the gel as described (PINTO *et al*. 1994) and were purified by ethanol precipitation. To generate the first strand cDNA, 5′-adaptor-ligated products were mixed with 0.3 μl of 100 μM s1082 oligonucleotide containing a randomized 9 nt sequence, incubated at 65°C for 5 min, and cooled to 4°C. A mixture of 4 μl of 5× First-Strand buffer, 1 μl (40 U) RNaseOUT, 1 μl of 10 mM dNTP mix, 1 μl of 100mM DTT, 1 μl (200 U) of SuperScript III Reverse Transcriptase and 1.7 μl of nuclease-free water was added to the reaction. The reaction was incubated at 25°C for 5 min, 55°C for 60 min, 70°C for 15 min, and then cooled to 25°C. 10 U RNase H was added, the mixtures were again incubated for 20 min at 37°C, and 20 μl of 2× DNA loading solution (PippinPrep Reagent Kit, Sage Science) was added. Then, the DNAs were separated by electrophoresis on 2% agarose gel (PippinPrep Reagent Kit, external Marker B) to collect a library of ∼90 to ∼ 550 nt range which is recovered by ethanol precipitation. To amplify cDNA, 9 μl of gel-isolated cDNA was added to the mixture containing 1× Phusion HF reaction buffer, 0.2 mM dNTPs, 0.25 μM of one of an i5 index containing lab-designed custom primer (CKO3414-3423), 0.25 μM of one of an Illumina TruSeq small RNA indexing primer (RPI1-32) and 0.02 U/μl Phusion HF polymerase in 30 μl reaction. Then, PCR was performed with an initial denaturation step of 10 s at 98°C, amplification for 12 cycles (denaturation for 5 s at 98°C, annealing for 15 s at 62°C and extension for 15 s at 72°C), and a final extension of 5 min at 72°C. Finally, PCR-amplified cDNA libraries were isolated by electrophoresis on 2% agarose gel and products in the range of ∼180 to ∼ 550 nt were collected following ethanol precipitation. Thus, generated dual indexed TSS-seq libraries which were sequenced together with STRIPE-seq libraries. TSS-seq and STRIPE-seq libraries were pooled and sequenced on an Illumina NovaSeq-S1-200 cycles sequencing flow cell using custom sequencing primer (s1115) and custom i5-index primer (CKO3424). **Supplement Figures S4a and S4b** show the library design schemes for both TSS-seq and STRIPE-seq.

### STRIPE-seq library preparation

Yeast cell cultures were grown in liquid YPD media and cells were harvested at the mid-log phase at a density of 1-1.5 × 107 cells/ml. STRIPE-seq library construction was performed as described by (POLICASTRO *et al*. 2020; POLICASTRO *et al*. 2021) with modifications on the oligos used, to make it compatible with the TSS-seq sequencing scheme. The steps are briefly described as follows; Total RNA was extracted, DNase I digested, and rRNA depleted as described above in the TSS-Seq method section. For a given library, two reactions apiece with 100ng rRNA-depleted RNA were set up in parallel. Terminator exonuclease (TEX) reaction, and template switching reverse (TSRT) reaction were performed as described in (POLICASTRO *et al*. 2020). For all the libraries, the same Reverse Transcription Oligo (RTO, CKO3425) with random nonamer was used instead of a unique RTO per library. See **Supplemental Table 3** for RTO and TSO sequences. Bead-purified TSRT products for two reactions per library were pooled into a single reaction before the final PCR. Final library amplification was performed on cleaned TSRT products using one of the i5 index containing lab-designed custom primers (CKO3414-3423), and one of the Illumina TruSeq small RNA index primers (RPI1-32) to generate dual indexed libraries. Amplified libraries were initially purified with SPRI select beads to remove small fragments and then gel extracted from 1.8% agarose gel to size select library products from 200-800 bp using NucleoSpin® Gel and PCR Clean-up (Macherey Nagel, NC0389463) kit. Hence, the generated STRIPE-seq libraries were sequenced with TSS-seq in the same Nova-Seq flow cell.

### TSS-seq data analysis

TSS-Seq data analysis followed the pipeline outlined in (ZHAO *et al*. 2021) (ARORA *et al*. 2025). Quality control, read trimming, and mapping were performed as described by (QIU *et al*. 2020; ZHAO *et al*. 2021), resulting in a TSS count table. Thus, the generated count table contains TSS-seq reads at each position within defined promoter windows (median TSS ± 250 nt upstream and 150 nt downstream) for 5,979 selected promoters. The data were then filtered to retain counts from n = X promoters with an average of ≥20 reads per promoter, which were used for downstream analysis. Key TSS metrics, such as ‘TSS spread’ and ‘TSS shift (or Median TSS shift), were calculated using scripts from (ZHAO *et al*. 2021). Differences in TSS spread and shift between WT and mutant for selected promoter classes were visualized as heatmaps (Morpheus) and/or boxplots.

### STRIPE-seq data analysis

The STRIPE-Seq data analysis pipeline was similar to TSS-Seq, generating metrics like ‘TSS spread’ and ‘TSS shift (or Median TSS shift).’ However, after alignment to the reference genome, the Picard toolkit removed reads with more than three soft-clipped bases. Downstream analysis from here was the same as described above in the TSS-seq data analysis section and in (ARORA *et al*. 2025).

## DATA AVAILABILITY

The sequencing data generated in this study for TSS-Seq and STRIPE-seq have been deposited in the NCBI BioProject database under accession number **PRJNA1346453**.

## ACKNOWLEDGEMENTS

We acknowledge funding from National Institutes of Health (NIH) grants R01GM097260, R01GM120450, and R35GM144116 awarded to CDK and for R35GM118059 awarded to BEN. We acknowledge funding from the National Science Foundation (US) to Texas A&M University for REU support for SEH (award DBI-0851611). For computing resources, we acknowledge funding to the University of Pittsburgh Center for Research Computing and Data (RRID:SCR_022735). Specifically, this award used the HTC cluster supported by NIH grant S10OD028483. We acknowledge Gabe Zentner and Robert Policastro for their STRIPE-seq protocol and Chhabi Govind for his valuable input on implementation of STRIPE-seq. We would like to dedicate this paper to the memory of Gabe Zentner.

---

**Figure S1:**
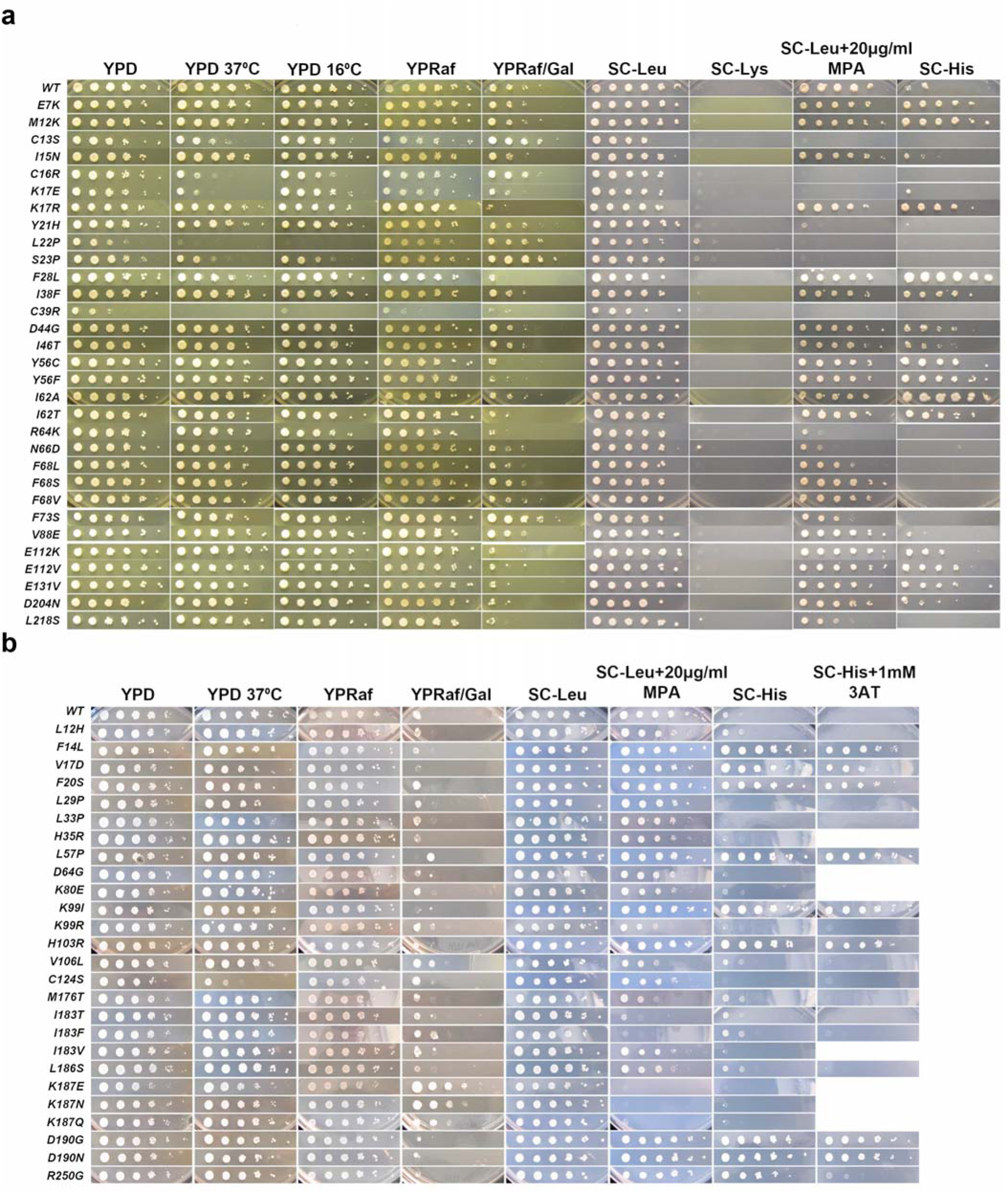
□Initiation□phenotypes of all□*tfb3*□and□*tfa1*□mutants obtained from genetic screening. □**a.**□Initiation□phenotypes of all□*tfb3*□mutants obtained from screening.□**b**.□Initiation□phenotypes of all□*tfa1*□mutants obtained from screening. Spot assays show the 10-fold serial dilutions of saturated cultures□of wild type and mutants plated on different□phenotyping□media.

**Figure□S2:**
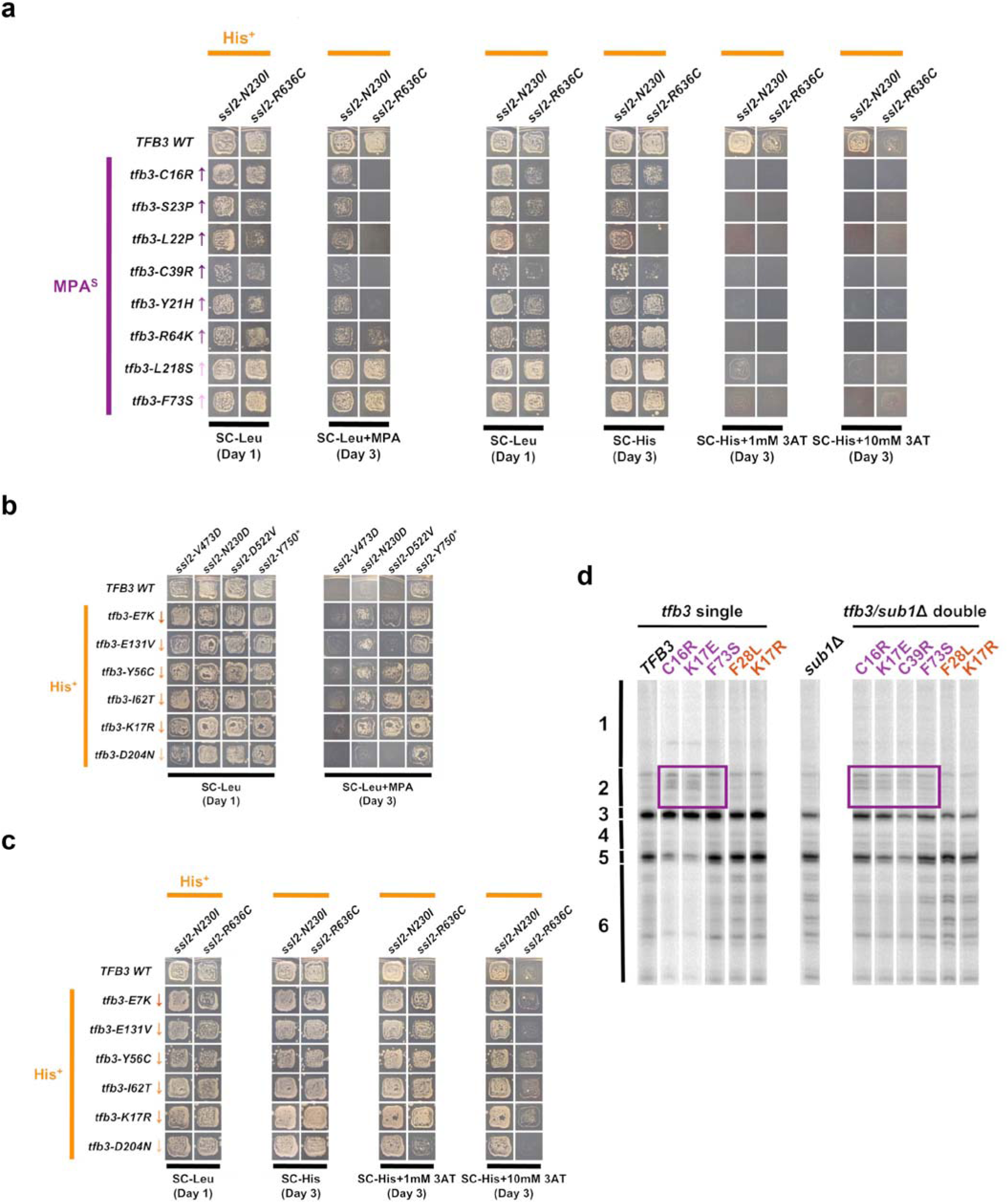
□Genetic interactions of□*tfb3*□alleges with□*ssl2*□and□*sub1*□show additive interactions. □**a.**□The patch phenotyping assay of□*tfb3*□MPA^S^/*ssl2*□His^+^□double mutants.□**b.**□Patch phenotyping assay of□*tfb3*□His^+^/*ssl2*□MPA^S^□double mutants.□**c.**□Patch□phenotyping assay of□*tfb3*□His^+^/*ssl2*□His^+^□double mutants.□**d.**□TSSs at□*ADH1*□detected by primer extension□for□*tfb3*,□*sub1*Δ,□*and tfb3/sub1*Δ□mutants.

**Figure S3:**
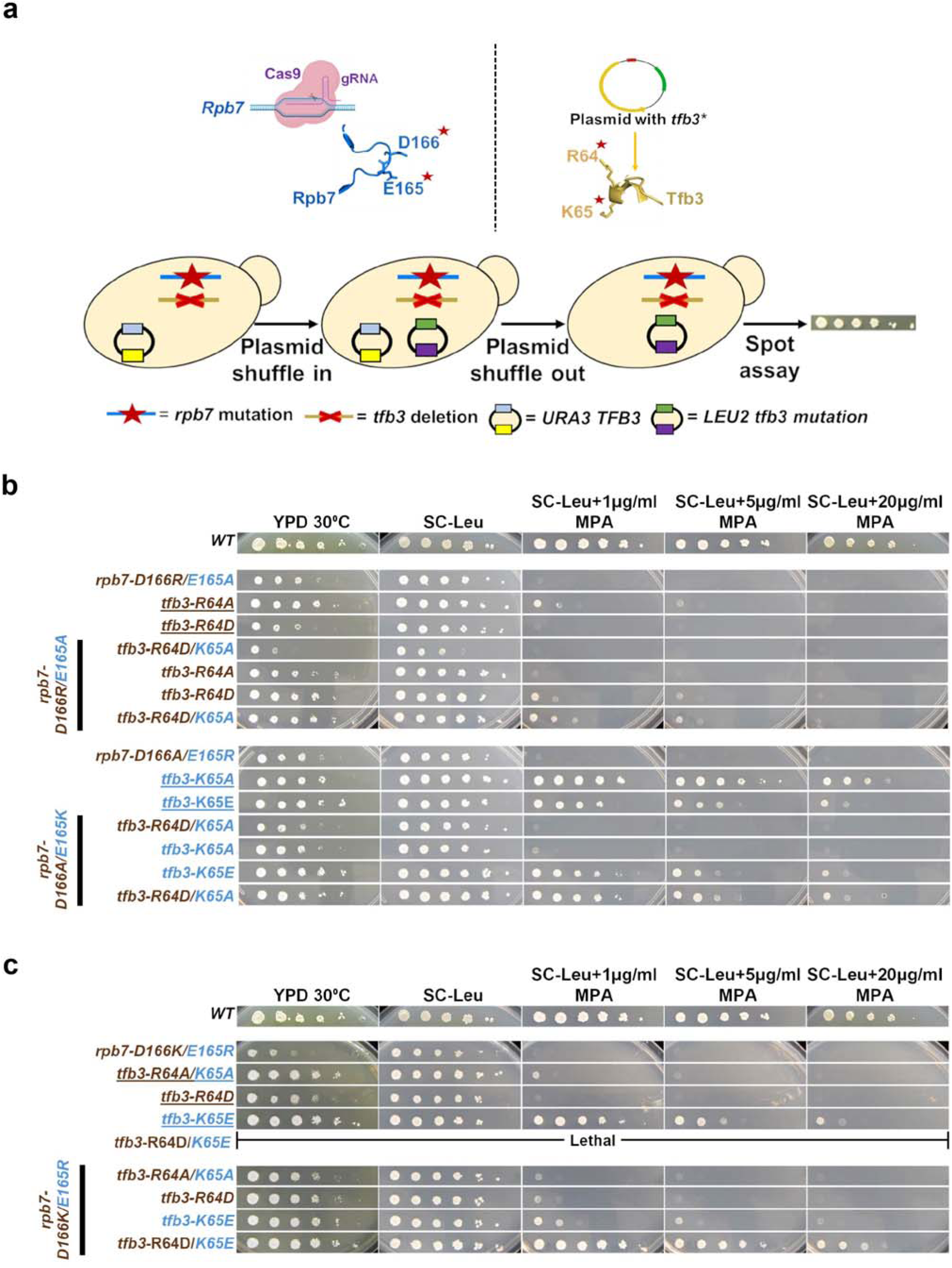
□Additional double mutant analyses provide evidence for the functional coupling between both salt bridge-forming residue pairs. **a**. Schematic representation of CRISPR editing and PCR sewing followed by plasmid shuffle to generate mutants at salt bridge-forming residues.□**b.**□The charge reversal at individual salt bridges partially suppresses double□SB mutants.□**c.**□An alternative□*rpb7*□mutant□i.e.□*rpb7-D166K/E165R + tfb3-R64D/K65E*□also shows□suppression□through□charge□reversal.

**Figure S4:**
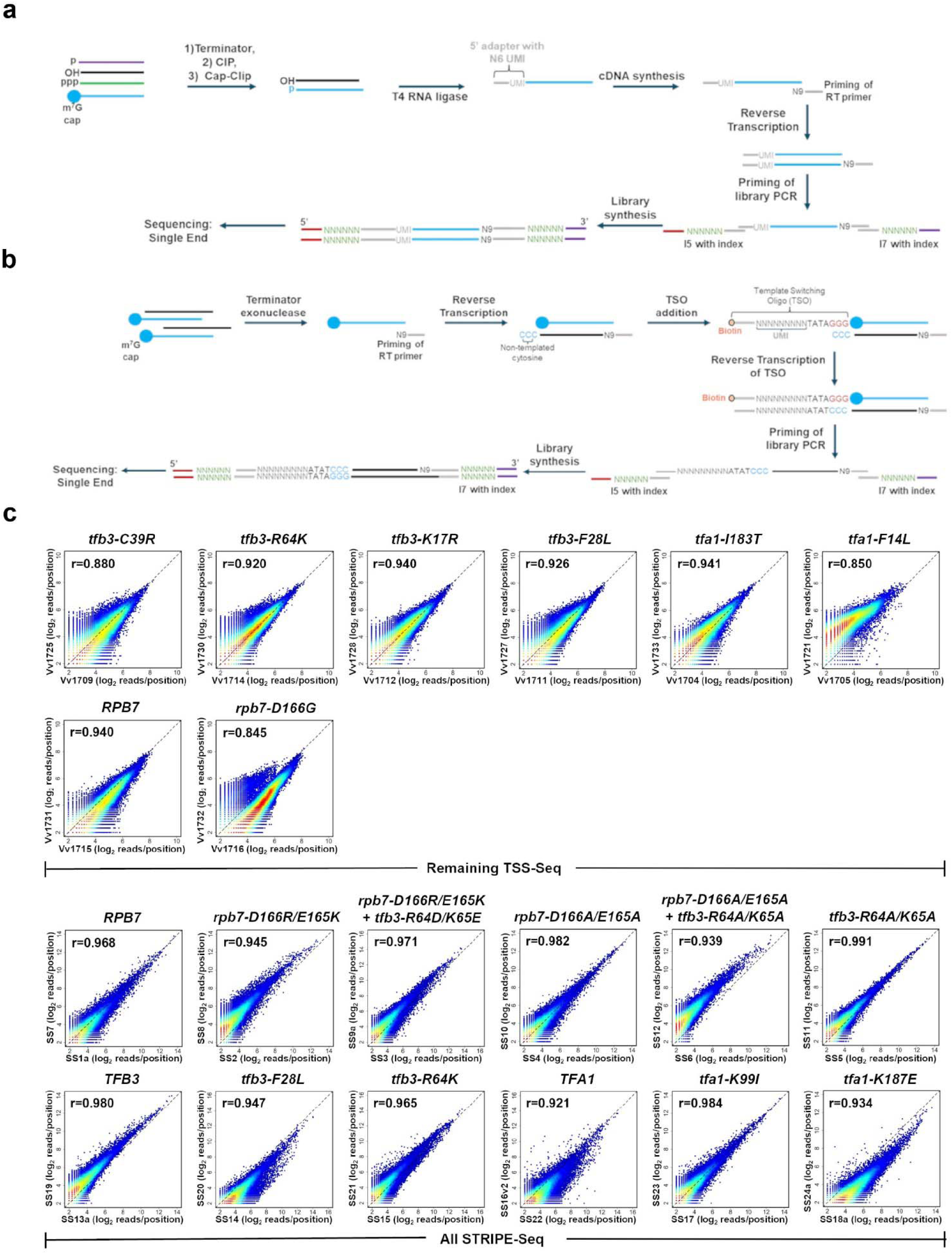
The transcription□start□site sequencing (TSS-seq) and STRIPE-Seq□identify□global initiation effects. **a.**□TSS-seq library construction is based on the protocol described by Vvedenskaya et al., 2015 and Qiu et al. 2020, with dual-index oligos replacing the single-indexed approach during the final library amplification step (Vvedenskaya *et al*. 2015; Qiu *et al*. 2020), see Materials and Methods. **b.** STRIPE-seq library construction is based on the protocol described by Policastro et al., 2020 (Policastro *et al*. 2020), implementing a single unique reverse transcription oligo (RTO) strategy instead of a unique RTO per library, see Materials and Methods. **c.** Scatter plot showing the correlation of log_2_ transformed reads at individual genome positions for all positions ≥ 3 reads for biological replicates of TSS-Seq and STRIPE-Seq libraries.

**Figure S5:**
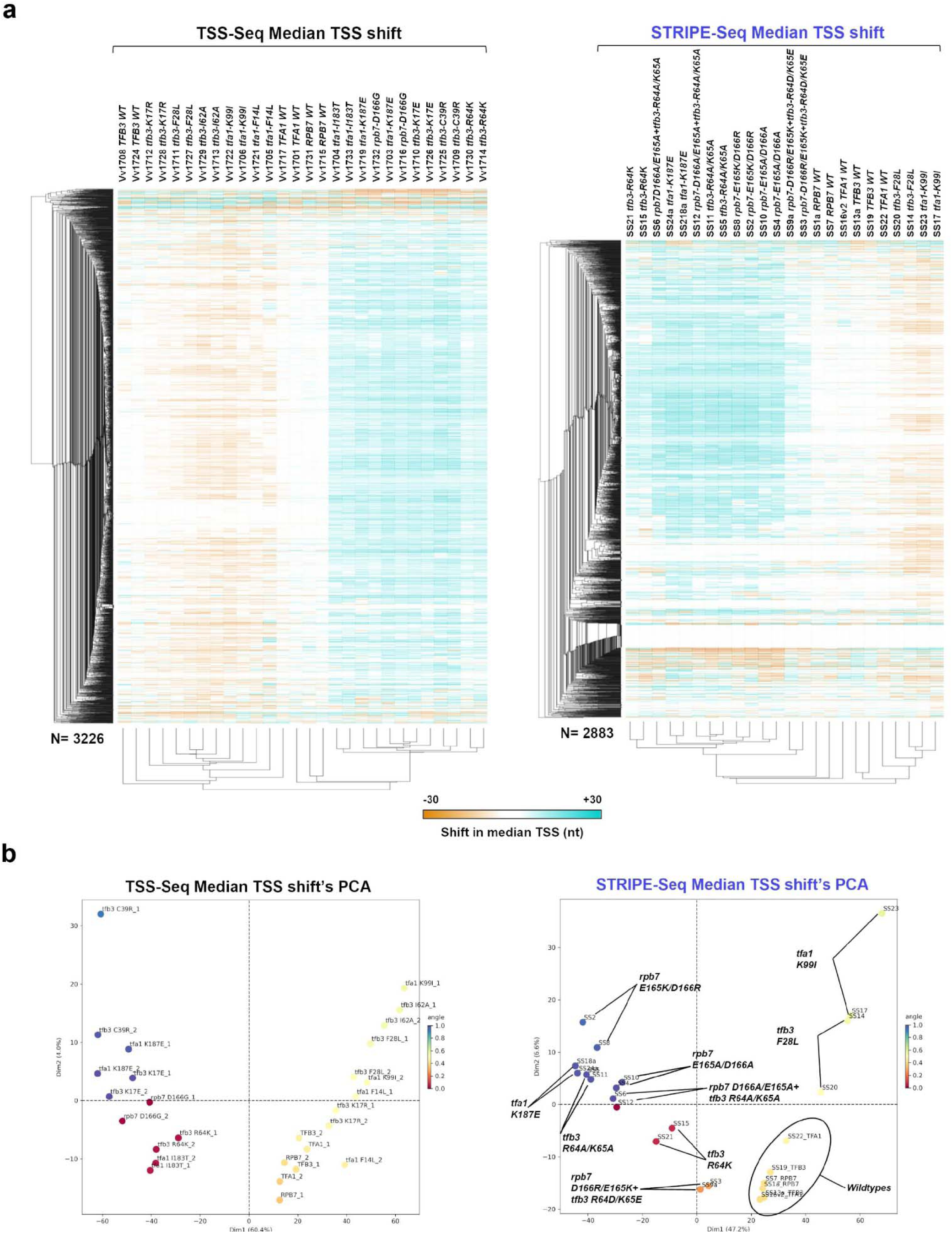
Analysis of transcription start site (TSS) shifts in mutants via TSS-Seq and STRIPE-Seq. **a.**□Heatmap showing hierarchical clustering of median TSS shifts for□*tfb3*,□*tfa1*, and□*rpb7*□mutants across individual replicates (TSS-Seq; left and STRIPE-Seq; right). Data is presented as in□**Figure**□**6d**, but for individual replicate libraries.□**b.**□The principal□component□analysis (PCA) for median TSS shift□*tfb3*,□*tfa1*, and□*rpb7*□mutants in individual replicates (TSS-Seq; left and STRIPE-Seq; right).

**Figure S6:**
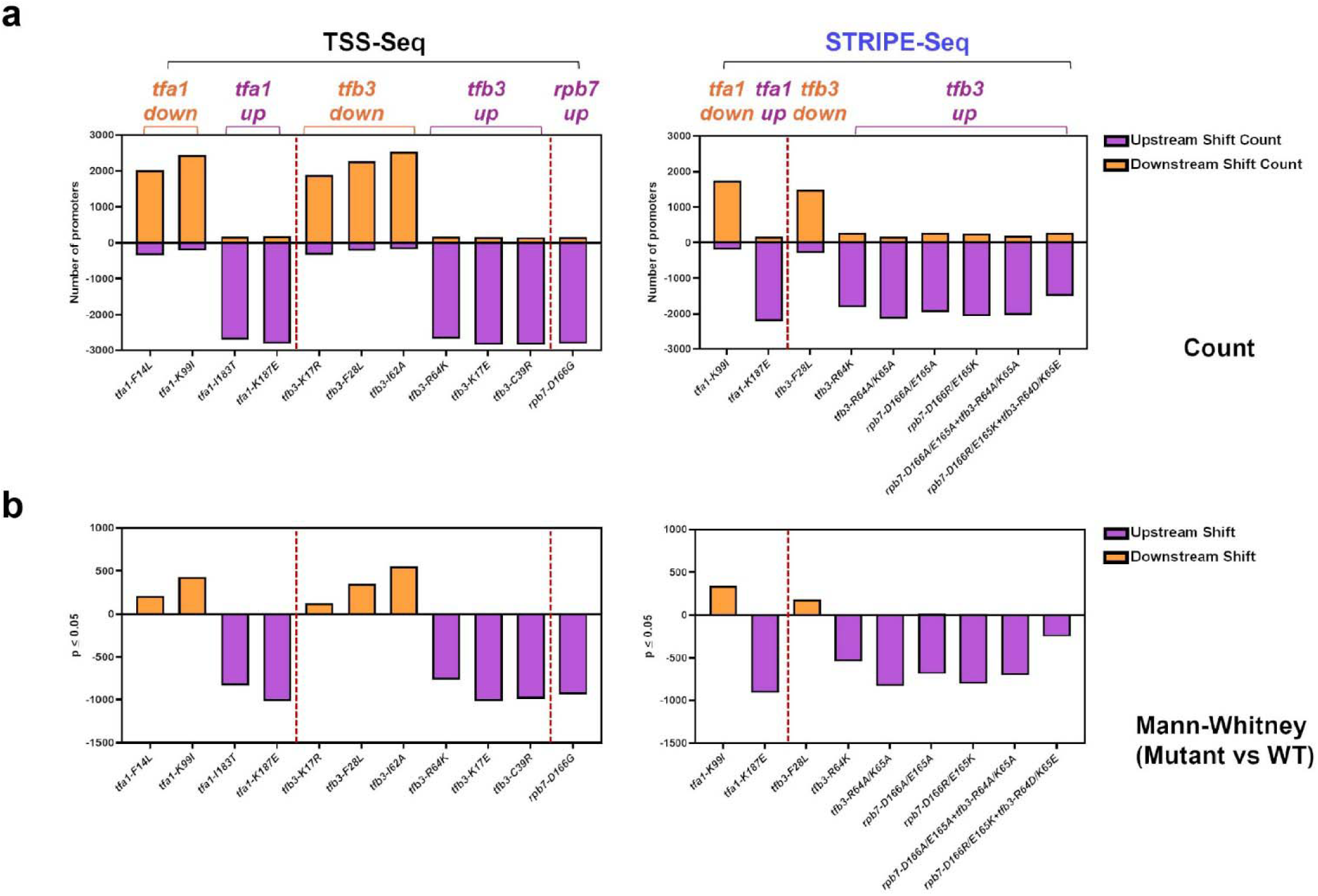
Promoter-level effects of TSS-shifting mutants on transcription start site selection. **a.**□The direction of TSS shift depends on the given mutant class. Counts of promoters with upstream shifts are shown below the x-axis (negative numbers) and counts of promoters with downstream shifts are shown above the x-axis. The mutants analyzed by TSS-Seq are shown on the left, and those analyzed by STRIPE-Seq are shown on the right. **b**. Statistical analysis of significant TSS shifts at individual promoters. For each mutant, TSS shifts at individual promoters across biological replicates were compared with those observed for WT. Promoter-specific TSS shifts were calculated relative to the median WT TSS position determined from aggregated WT data. Numbers shown indicate the count of promoters with significant upstream (negative) or downstream (positive) TSS shifts, as determined by the Mann–Whitney U test (p ≤ 0.05). Statistical tests were performed independently for each promoter using TSS shift values from all strains (n = 2 for mutants and n = 6 for WT). The mutants analyzed by TSS-Seq are shown on the left, and those analyzed by STRIPE-Seq are shown on the right.

**Figure S7:**
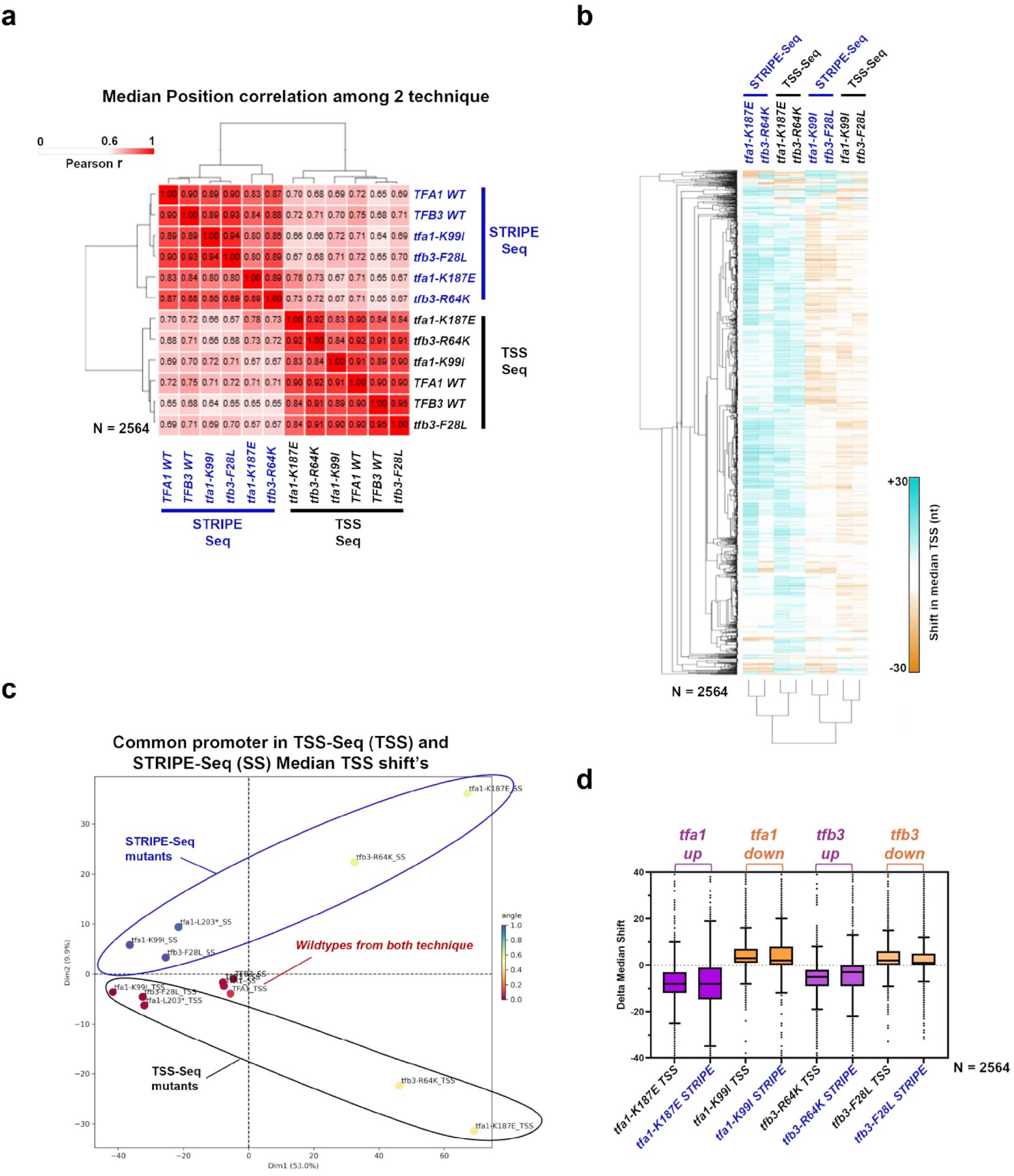
Mutant behavior seems consistent between TSS vs STRIPE-seq techniques with technique driven variability. **a.**□Hierarchical clustering of Pearson correlation coefficients for□*tfb3*, and□*tfa1*□median shift position in 2654 promoters common to both techniques.□**b**. Heatmap showing hierarchical clustering of median TSS shifts for□*tfb3*, and□*tfa1*□mutants (promoters as in (a)).□**c.**□The principal□component□analysis (PCA) for median TSS shift in□*tfb3*, and□*tfa1*□mutants shows variation among mutants is technique driven.□**d.**□Box plot showing median TSS shifts across promoters in *tfb3*, and *tfa1* mutants. Techniques differ in magnitude at promoters but consistently capture median TSS shift at the individual promoter level. Blue labels for libraries that are sequenced by STRIPE-Seq and black for TSS-Seq. Median TSS shifts across promoters, are statistically distinguished from wildtype at p < 0.0001 for all mutants (Wilcoxon signed rank test).□

